# Immediate early gene expression dynamics in vivo segregates neuronal ensemble of multiple memories

**DOI:** 10.1101/2020.12.17.423270

**Authors:** P. Meenakshi, S. Kumar, J. Balaji

## Abstract

Immediate early genes (IEGs) are widely used as a marker for neuronal plasticity. Here, we model the dynamics of IEG expression as a consecutive, irreversible first order reaction with a limiting substrate. We show that such a model, together with two-photon in vivo imaging of IEG expression, can be used to identify distinct neuronal subsets representing multiple memories. We image retrosplenial cortex (RSc) of cFOS-GFP transgenic mice to follow the dynamics of cellular changes resulting from both seizure and contextual fear conditioning behaviour. The analytical expression allowed us to segregate the neurons based on their temporal response to one specific behavioural event, thereby improving the sensitivity of detecting plasticity related neurons. This enables us to establish representation of context in RSc at the cellular scale following memory acquisition. Thus, we obtain a general method which distinguishes neurons that took part in multiple temporally separated events, by measuring fluorescence from individual neurons in live mice.

**Summary:** Identifying neuronal ensemble associated with different memories is vital in modern neuroscience. Meenakshi et al model and use the temporal expression dynamics of IEGs rather than thresholded intensities of the probes to identify the neurons encoding different memory in vivo.

**Graphical abstract:** 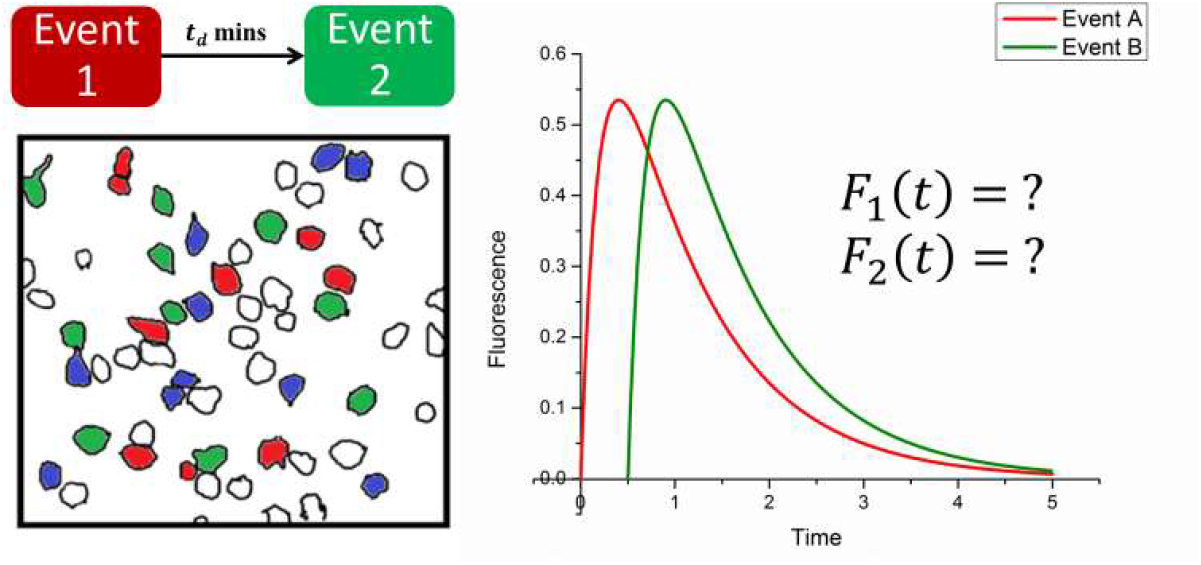

## Introduction

Identifying the neuronal ensemble that possibly encode memory and understand the changes happening in the memory circuits overtime are pertinent problems in neuroscience research(Cajal, 1888)(Josselyn et al., 2015)(Richards and Frankland, 2017)(Josselyn and Tonegawa, 2020). Immediate early genes are rapidly and transiently expressed in response to neuronal activity and hence used as a marker for plasticity. Generally, using IEG based techniques to identify the temporal coupling of IEG expression to different behaviours or events requires as many distinct molecular labels as the number of events that are being followed.

However, cellular compartment analysis of temporal activity by fluorescent in situ hybridization (catFISH)(Guzowski, 2002a; Guzowski et al., 1999; Guzowski and Worley, 2001) exploits the unique transport kinetics of Arc mRNA from the nucleus to the cytoplasm but the method is limited to in vitro identification. Another method uses a combination of IEG promoter-based expression and a modified Tet-OFF system to achieve labelling of two distinct population of neurons(Reijmers et al., 2007) (ref: Mayford,). In both cases, the visualisation of the signal is done post hoc in vitro, limiting the investigation of the neuronal population to a snapshot at any given time. Such methods cannot follow an ensemble of neurons and observe their evolution longitudinally.

Neuronal activity indicators such as GECIs or GEVIs are reporters of neuronal firing in sub millisecond time scales(Lin and Schnitzer, 2016; Knöpfel, 2012). However, they do not necessarily report the plastic events that occur in response to these firings. Further, they require high speed imaging with temporal resolution matching the response time of the indicators often requiring the imaging be done on awake behaving mice. Slow kinetics of IEG expression enables, following cellular plastic events in anesthetised mice after behavioral training. Thus, in IEG based methods the likelihood of the active neuron to be plastic and more importantly it makes it easier to combine with conventional behavioral paradigms to study memory.

Here, we propose to follow the kinetics of IEG expression *in vivo* to identify distinct neuronal subsets each corresponding to distinct events. cFOS expression is one of the widely used marker for cellular activity (Dragunow and Faull, 1989). First, we analytically describe the IEG protein expression dynamics as a function of time. In case of a fluorophore expressed under an IEG promoter, the kinetics can be followed by measuring the fluorescence as a function of time. Next, we validated the model using data obtained from cfos-egfp (Barth et al., 2004a) and cfos-shgfp (Reijmers et al., 2007) transgenic mice under two conditions: bicuculline induced seizure and behavioural induction of IEG following contextual exposure. Finally, the kinetics of IEG-fluorescence expression was used to estimate when the activity of the neuron was induced, enabling us to distinguish between neuronal populations that took part in different events that are separated in time. Thus, we hypothesize that the memory engram of multiple events can be identified using the expression dynamics of an immediate early gene.

Recent and emerging evidence suggest RSc plays a vital role in encoding context related information(Mitchell et al., 2018; Auger and Maguire, 2018; Auger et al., 2012, 2015; Miller et al., 2014). Clustered addition of spines is observed in RSc when contextual training is carried out across multiple sessions(Frank et al., 2018). Similarly, inactivation of RSc prevents contextual retrieval in mice both after contextual fear conditioning and water maze training(Opalka and Wang, 2020)(Czajkowski et al., 2014). Preferential activation of RSc during spatial navigation has also been reported (Cowansage et al., 2014). Interestingly, it is also shown that in schema dependent encoding of related events, RSc is engaged only during the encoding of new learning related to prior information but not during encoding of completely novel information(Tse et al., 2011). More importantly, RSc lesion in rats abolishes its ability to resolve context-based conflicts (Nelson et al., 2014). Thus, all these studies suggest RSc plays an active role during the context based behavior, although it is not clear what is the nature of this role. Given its role, we reasoned that it is possible RSc might maintain contextual information in form of independent representations and its interrelations. If such contextual interrelations, have cellular representations in RSc, we would be able to locate them using our method. This is possible as the method described here, can identify, and longitudinally follow the activated cellular ensembles in vivo. Thus, in this study we simultaneously probe representation for a context, how it changes across time, and when a new context is introduced close in time.

## Theory

### Analytical description of IEG expression dynamics

In order to model IEG expression kinetics in a neuron, we assumed that a pool of mRNA is present which is translated to protein in response to behaviourally relevant neuronal activity or signal (Greenberg et al., 1986)(Saha et al., 2011). In response to a signal, mRNA (A) is converted to protein (B) with a rate constant of protein formation (k_f_). One of the marked features of IEG’s is their transient nature. As the protein forms, ubiquitination degrades the protein, and we assume such a reaction to be first order in protein concentration with a degradation rate constant of (k_d_). Thus, the IEG expression kinetics can be considered as consecutive first-order reactions (Fig1(a)).

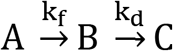

where A is the concentration of the substrate for protein synthesis, B represents the number of protein molecules and C is that of the degraded products.

**Figure 1:**
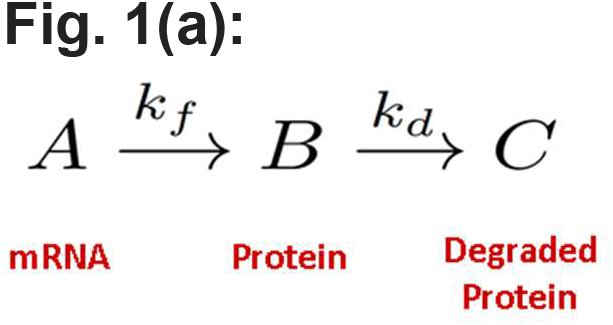
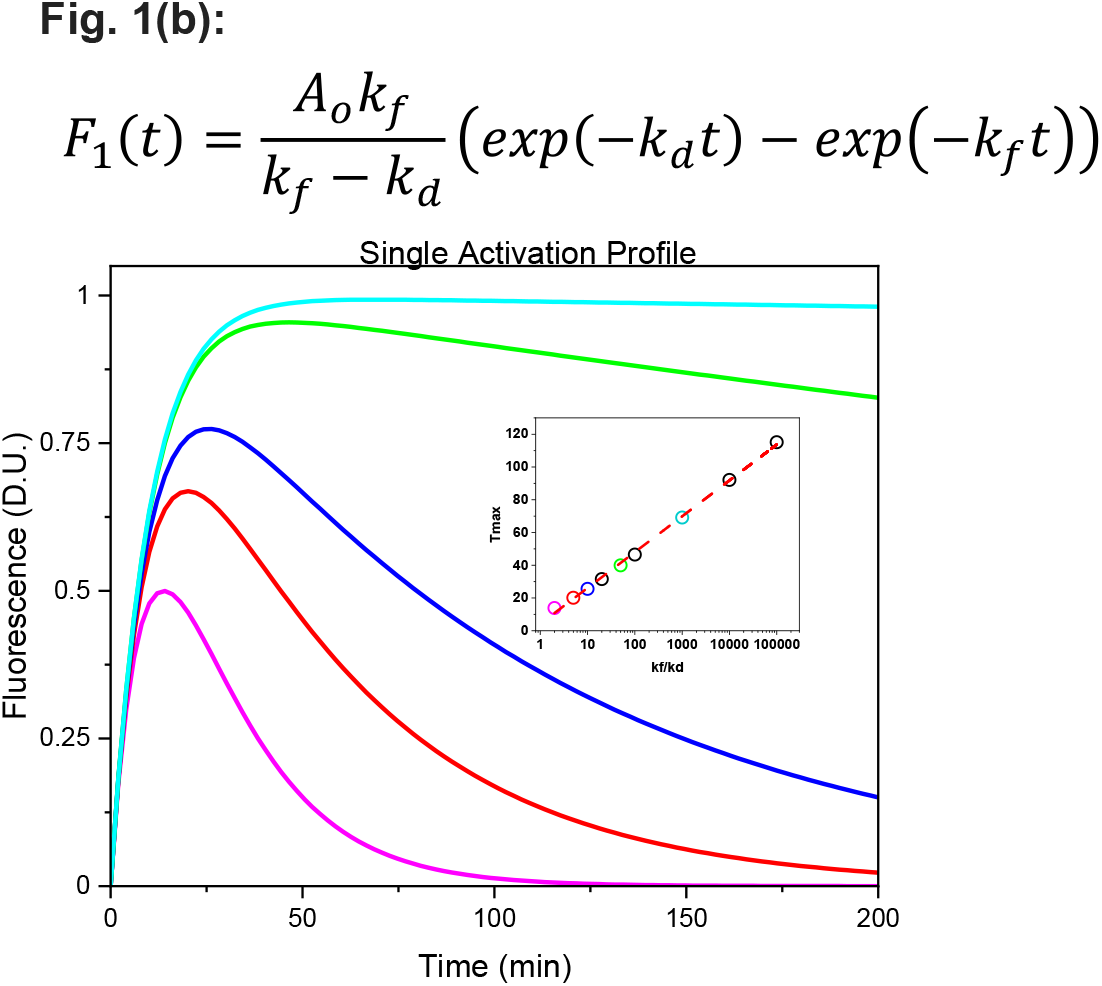
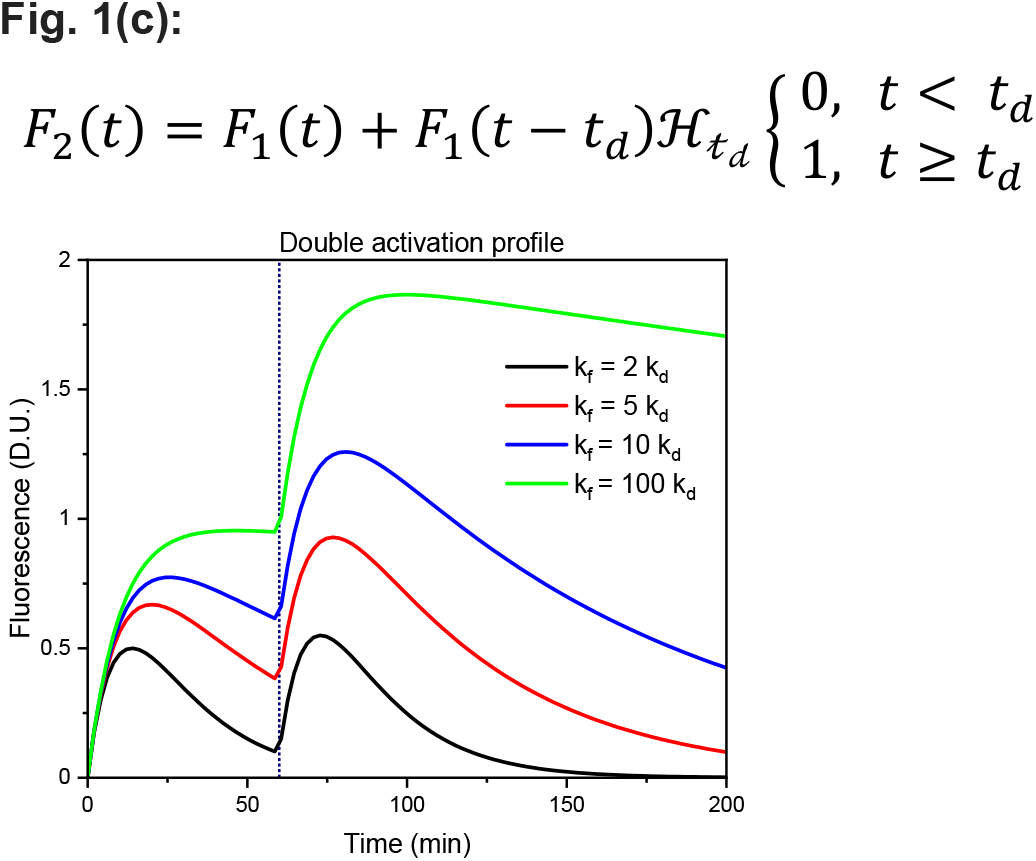
Analytical description of an IEG expression in response to plasticity related events. (a) A simple consecutive reaction kinetics for the mRNA (A), protein (B) and degraded protein (C) describes the response to a plasticity signal. The reaction is assumed to proceed with first order forward reaction rate constants *k_f_* and *k_d_* for the synthesis and degradation of proteins, respectively. (b) Solving the coupled differential equations of the rate equations of (a), we get an expression (Eq. 1) for F1(t) that describes the fluorescence intensity corresponding to IEG coupled fluorophore at any given time ‘t’ as a difference of two exponential terms with rate constants, *k_f_* and *k_d_*. The lines are the simulated response functions for five values of k_f_/k_d_ ratios with parameter A and k_f_ set to 1 D.U and 0.1 min^-1^). Time to maximal response, one of the key parameters necessary to time the neuronal tagging is plotted for these set of ratios as a scatter plot in the inset. The color of the open circles correspond to their respective solid lines. The red dashed line is a straight line fit of these scatter plot. (c) Similarly we describe Eq. 2 for a neuron that got activated twice where A, k_f_, k_d_ are as previously described and t_d_ is the time of second activation event. Eq. 2 is simulated (solid lines) to show the response for four ratios of *k_f_/k_d_* with parameter A, k_f_ and k_d_ set to 1D.U., 0.1min^-1^and the time gap between the two events (t_d_) is set to 60 min as indicated by the black dotted line.

Given the first order nature of the reaction we proceed to write the rate equations for A and B as follows,

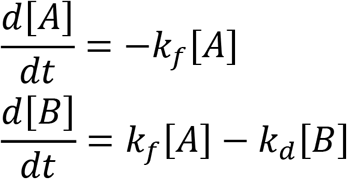

In order, to solve for the [B] the number of protein molecules that is going to be present transiently we need to solve the above coupled differential equations. We proceed by solving for B in Laplace space. Thus, using Laplace transform to transform the equations above we get,

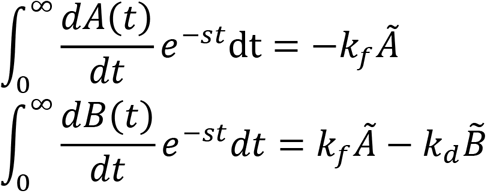

which can be reduced to the below equations,

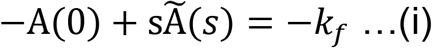

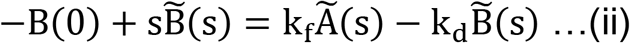

where A(0) is the maximum substrate concentration per signalling event. B(0) is the concentration of B at time t = 0, hence B(0) = 0. Rearranging equation (i), we get the value of A(s).

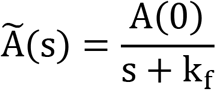

Using this in equation(ii) and rearranging for ~B(s), we get,

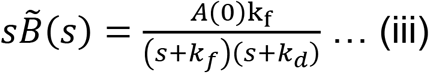

Equation (iii) represents the number of IEG molecules in Laplace space. Using partial fractions and obtaining the inverse Laplace transform, we have,

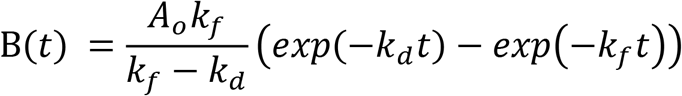

A fluorescence signal from a sample (F(t)) is proportional to the number of molecules (B(t)). Using the quantum efficiency (ϕ_f_), the absorption cross-sectional of EGFP (∈_A_) and the collection efficiency (C_f_). we can write,

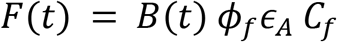

For a given imaging system, the fluorescence signal (F(t)) is directly proportional to the number of molecules present at any given time (B(t)).

Thus, the equation can be rewritten as,

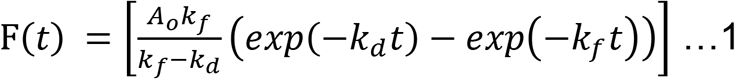

Using a similar approach, we can generalise the equation for multiple activation events in a neuron, as follows:

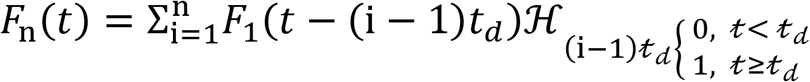

In case of a double activation event, the equation simplifies to:

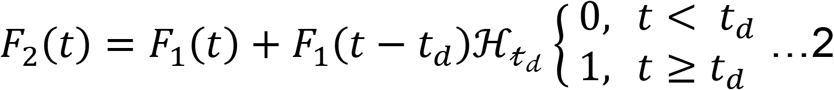

where t_d_ is the time of the second activation event or signal.

## Results and Discussion

### A consecutive first-order kinetics describes IEG protein expression following bicucline administration in cfos-egfp transgenic mice

In order to investigate whether the derived analytical expression (see: Theory) describes IEG protein expression *in vivo*, we imaged cFOS expression in response to administration of bicuculine. Bicuculine is known to induce seizures and cFOS expression in mice(Morgan et al., 1987)). The mice we used expresses the transgene cFOS EGFP in response to cFOS induction. Immediately after the seizure, mice were subjected to *in vivo* imaging of the RSc (Fig 2(a)). Imaging time points ranged from 20-280 mins, that are ~10 minutes apart, keeping the imaging setup parameters viz the incident power (30 mW), pulse width (~100 fs), excitation wavelength (900 nm) and gain of the detection system constant. The fluorescence measured from individual nucleus identified as circular ROI’s increased initially, reached a peak and then decayed to base line values. These values are then fit to the Eq. 1. We captured and utilised the vasculature of the brain as shown in Fig.4(a) to facilitate imaging of same region across days. These images together with the read out of x and y positions served as reference points. Fig 2(b) shows a representative snapshot of cfos-egfp fluorescence image of RSc in the mouse. The fluorescence response from few representative cells as a time series is presented in Fig 2(c). The fluorescence values from these cells were extracted as explained in Methods (Supplementary Figure S1). These fluorescence values are then used to obtain the cellular responses/activity as a time series. A total of 66 ROIs were identified for fluorescence extraction and their cellular activity profiles are obtained. These 66 ROIs were initially fit to eq1 to obtain the distribution of equation parameters A, k_f_, k_d_ (Supplementary Figure S2, Table 1). Figure 2(d) shows the cellular activity profile (open circles) of four representative cells and their fit (solid line) to Eq 1. The fit parameters are provided in Table 2. We see a good agreement of our model as measured by the Adj. R Sq > 0.5, in 80 % of the cells identified. We estimate the rise time of the fluorescence following activation and decay to be 1/k_f_ = 27 + 3 mins and 1/k_d_ = 200 + 28 mins (taking the faster component from Table 1) respectively from these fits. We used these rise and decay times to arrive at the sampling interval for further experiments.

**Figure 2:**
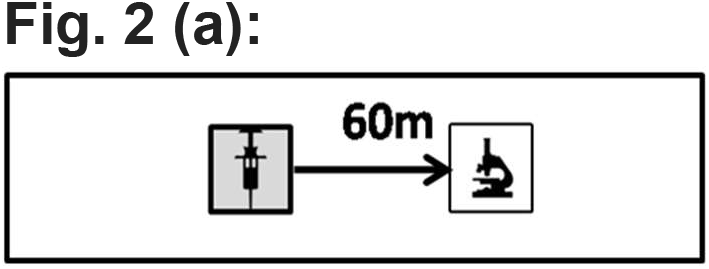
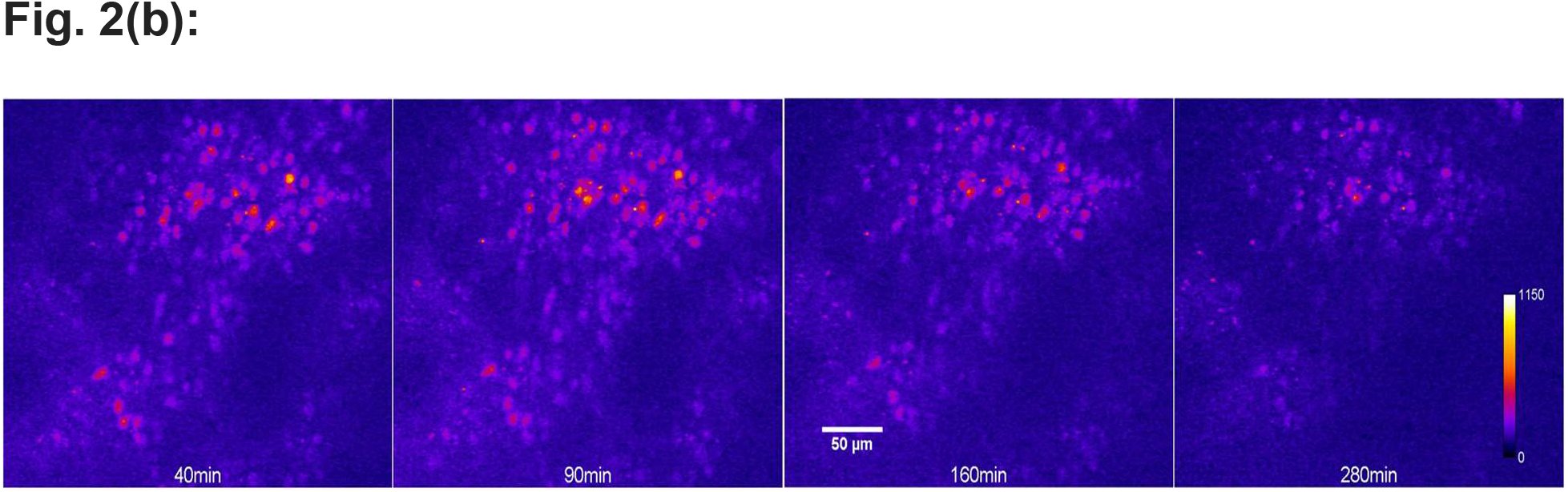
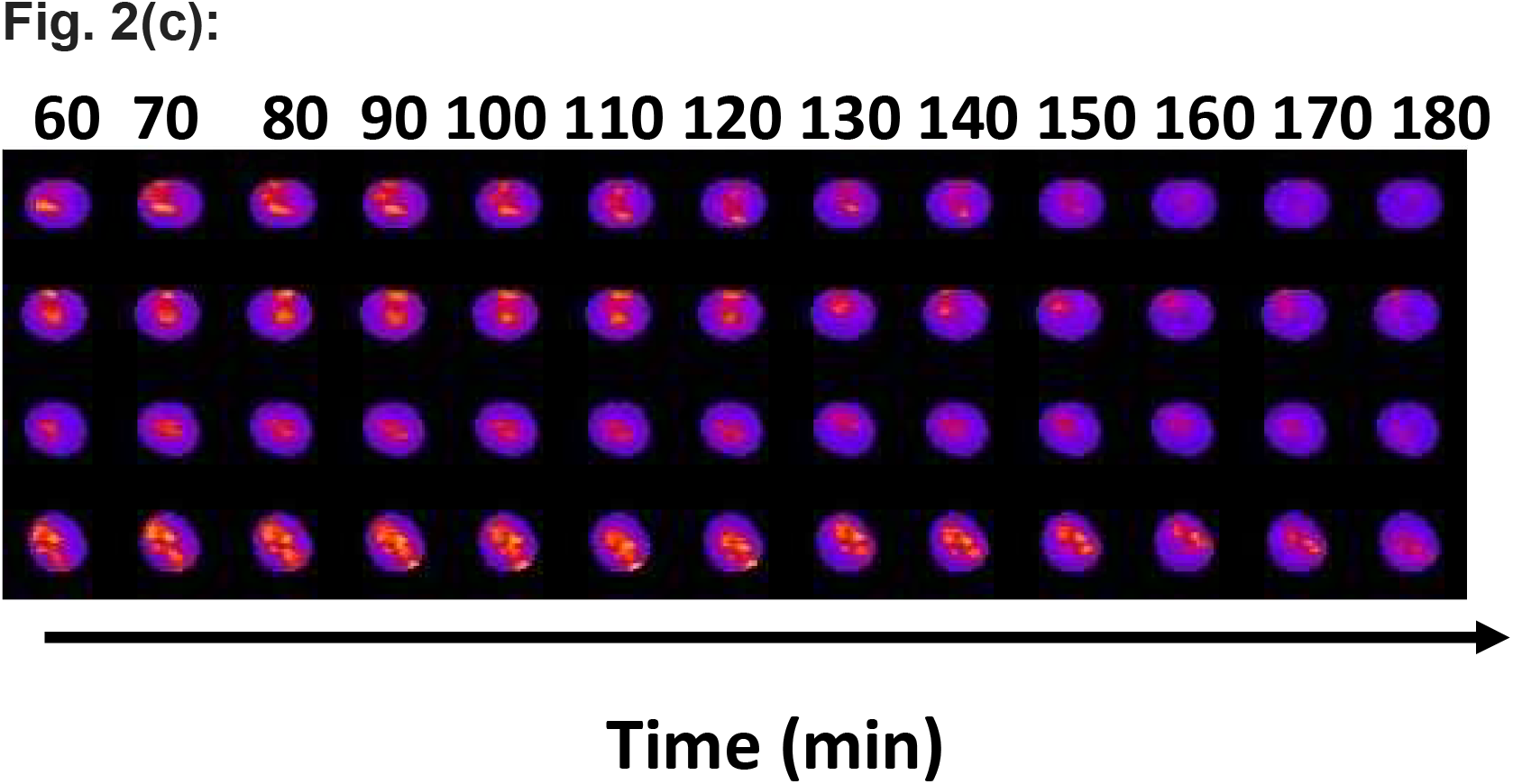
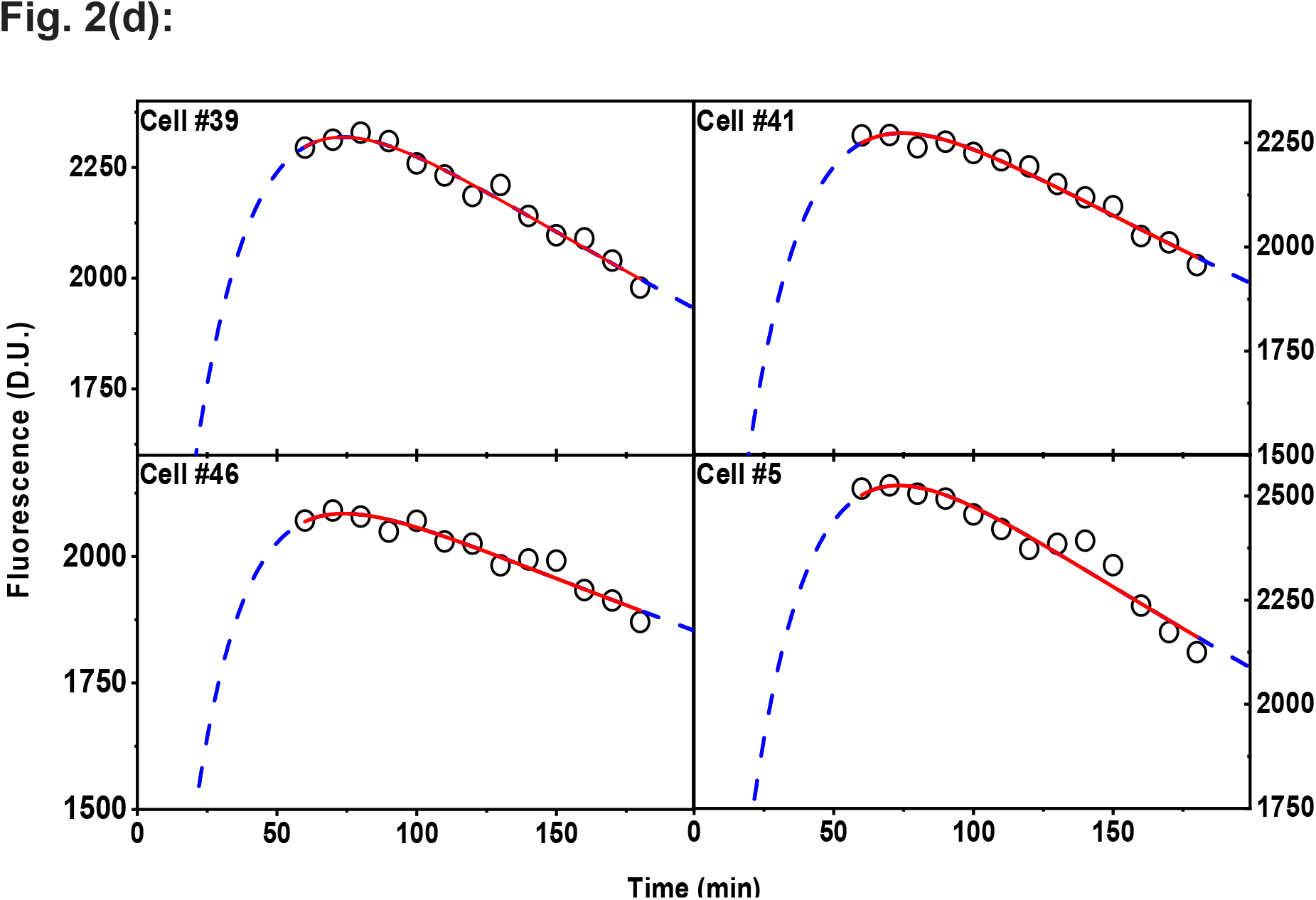
Seizure induced expression of IEG coupled fluorescence is described by first order consecutive kinetics: (a) Schematic of IEG induction to bicuculline administration and following its dynamics in an anesthetised transgenic mouse through in-vivo imaging of the retrosplenial cortex (RSc). (b) Maximum intensity projection of a stack of images corresponding to 200 x 200 x 200 *μ*m region obtained at different time points. The cFOS-EGFP signal is localised to the nucleus and hence the activated neurons appear as quasi circular regions of bright pixels with a diameter of ~20-pixel units. Snapshots RSc area shown are that of time points 40, 90, 160 and 280 mins. The scale bar in the image is X microns. (c) Select regions of interest centred around cells “#39”, “#41,” #46” and “#5”, are arranged as time series show the change in fluorescence across the entire cell nuclei. The scale bar in the image is 50 microns. Representative images of neurons in (d) across different time points. (d) Quantitative measure of cellular response extracted from the time series images through custom built software for four representative cells are shown as open circles. The open circles are obtained using the workflow (SFig. 1) and represent the activity of a neuron at a given time. The red line is the fit of this activity to Eq. 1. Blue dotted line extends the solid red line to the activity of the cell outside of the imaging time frame as predicted by our model. See table 1 for fit parameter details.

**Figure 3:**
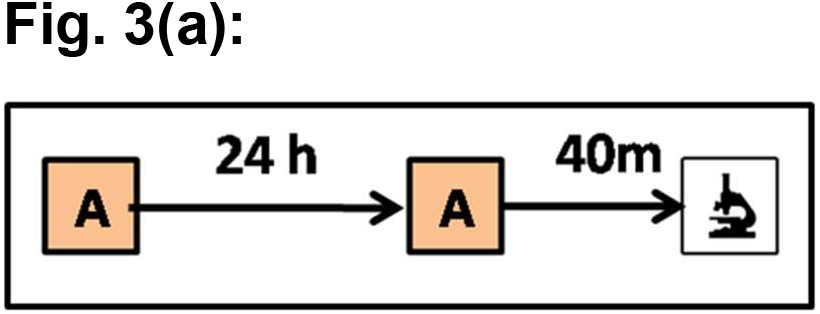
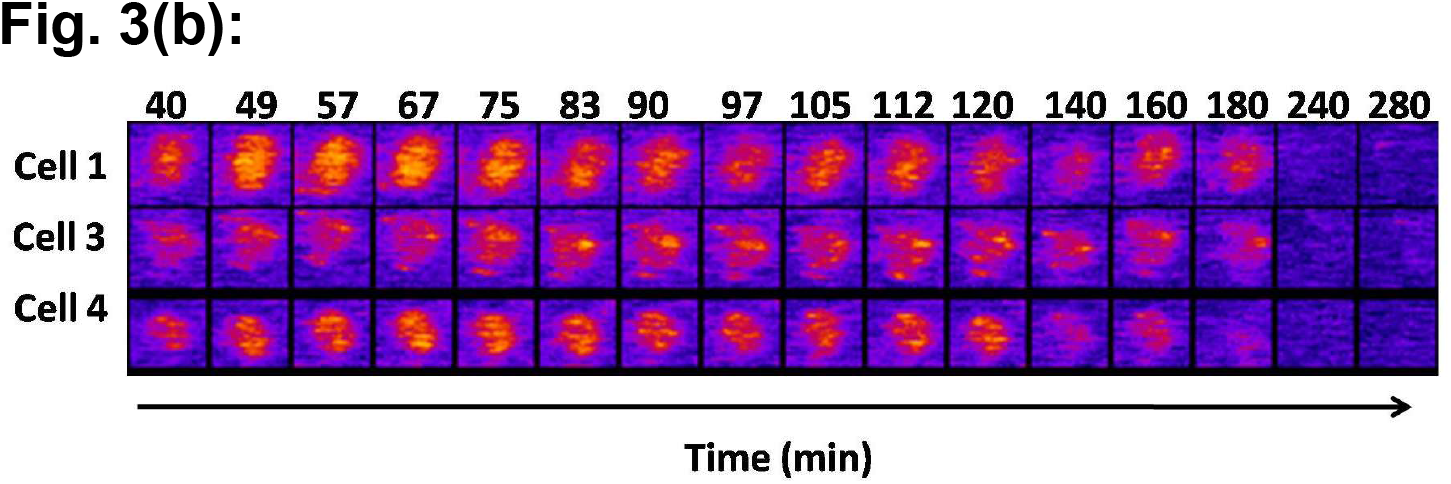
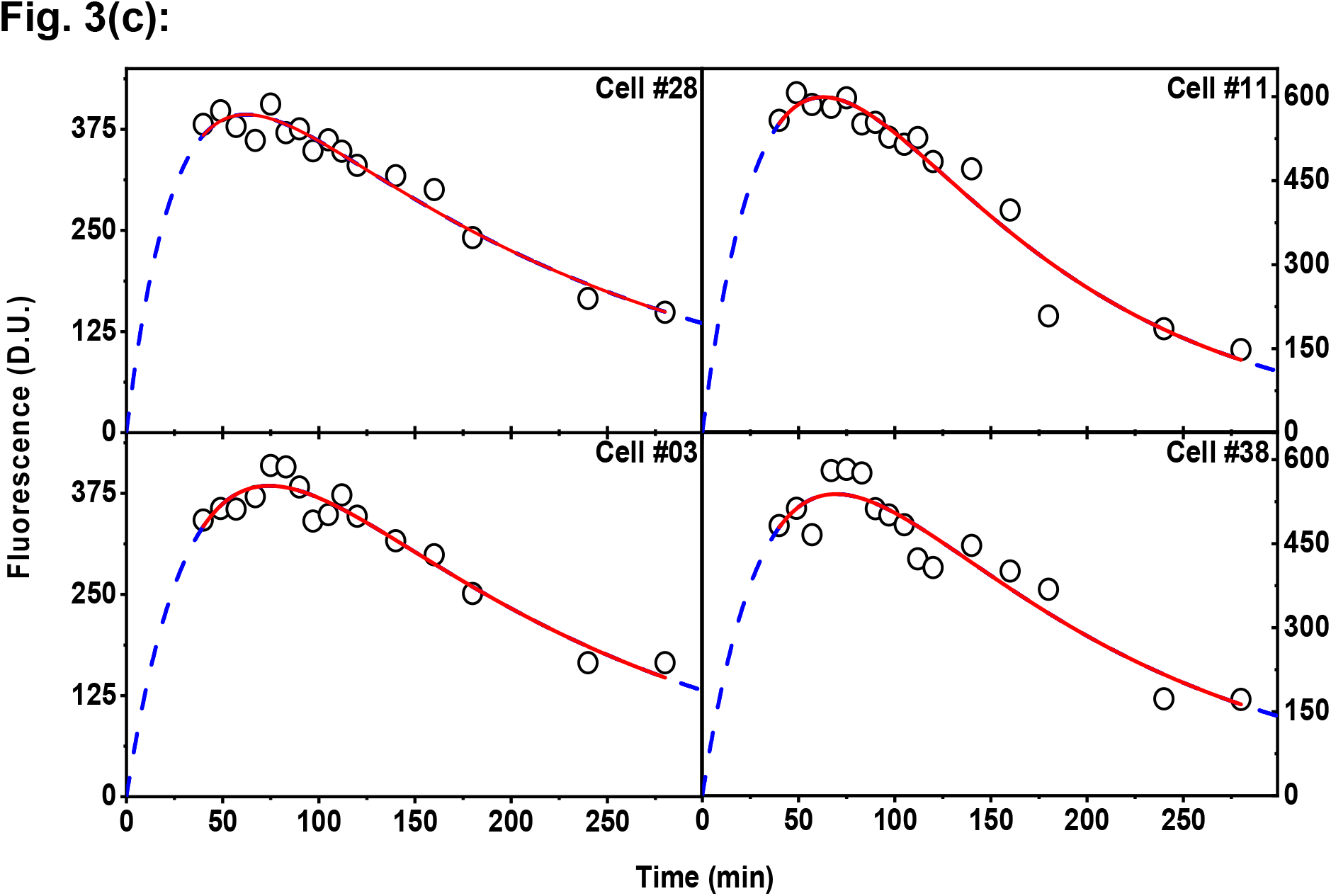
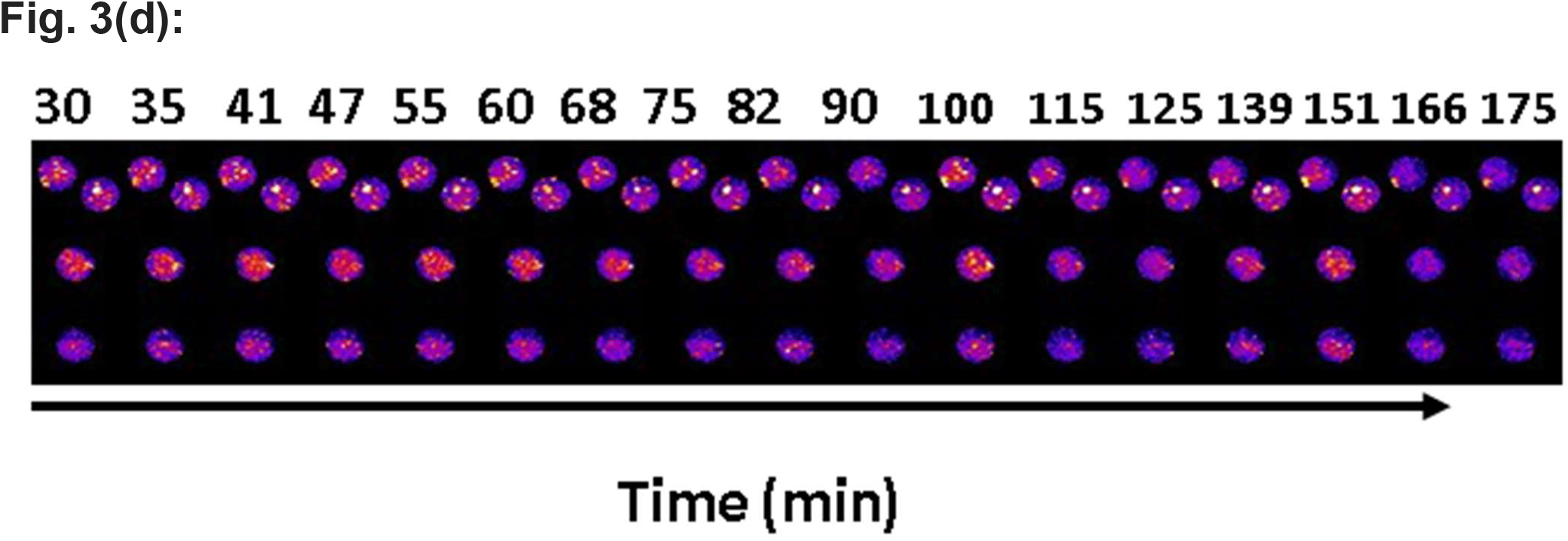
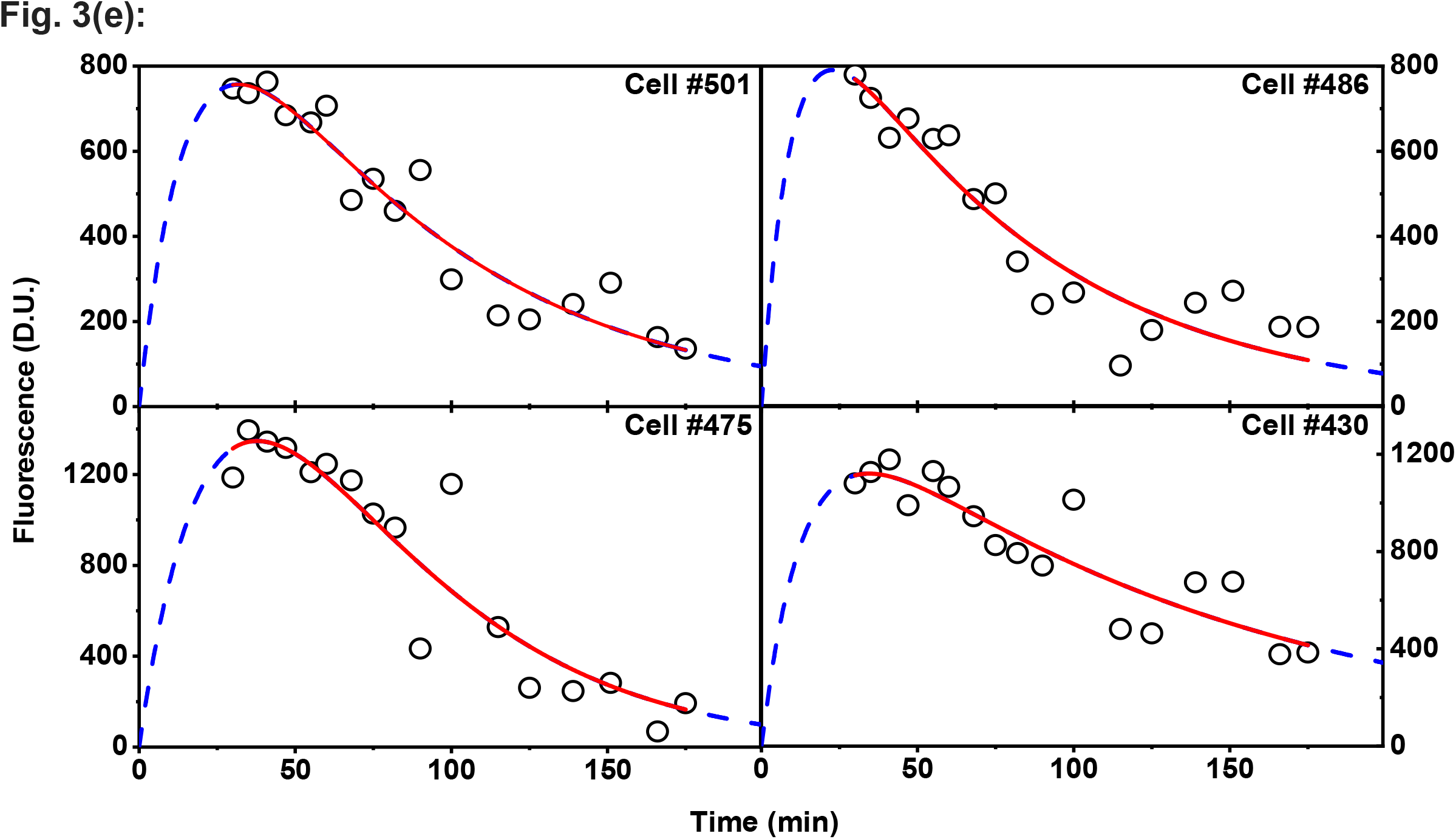
Quantification of behaviourally induced cfos-egfp and cfos-shgfp protein expression fits well to derived equation Fig. 1(b) indicating that the analytical equation generalises to different immediate early genes – fluorophore constructs. (a) Behavioral schematic used for inducing cFOS expression. Transgenic mice trained in context A and were made to recall the training context after 24 hours. The resulting activation of IEGs is followed immediately, through in-vivo imaging of the RSc in anesthetised mice. (b) and (d) are 4 representative image ROIs centred around cell “#501”, “#486”, “#475” and “#430” in cfos-egfp and cfos-shgfp transgenic mice across different time points respectively. (c) and (e) are the corresponding quantitative measure of cellular expression profile of cells in (b) and (d) along with their fits to Eq. 1. The open circles represent the amplitude of the cellular activity from a neuron at a given time. The red line is the fit of this data to Eq. 1. A good agreement of the fit to the observed data (Adj. R Sq > 0.92(cFOS eGFP), >0.79 (cFOS-shGFP)) indicates that our model is consistent with the observed cellular response. Blue dotted line extends the fits and spans the entire x-axis as explained in Fig. 2. See Table 1 for fit parameters.

**Figure 4:**
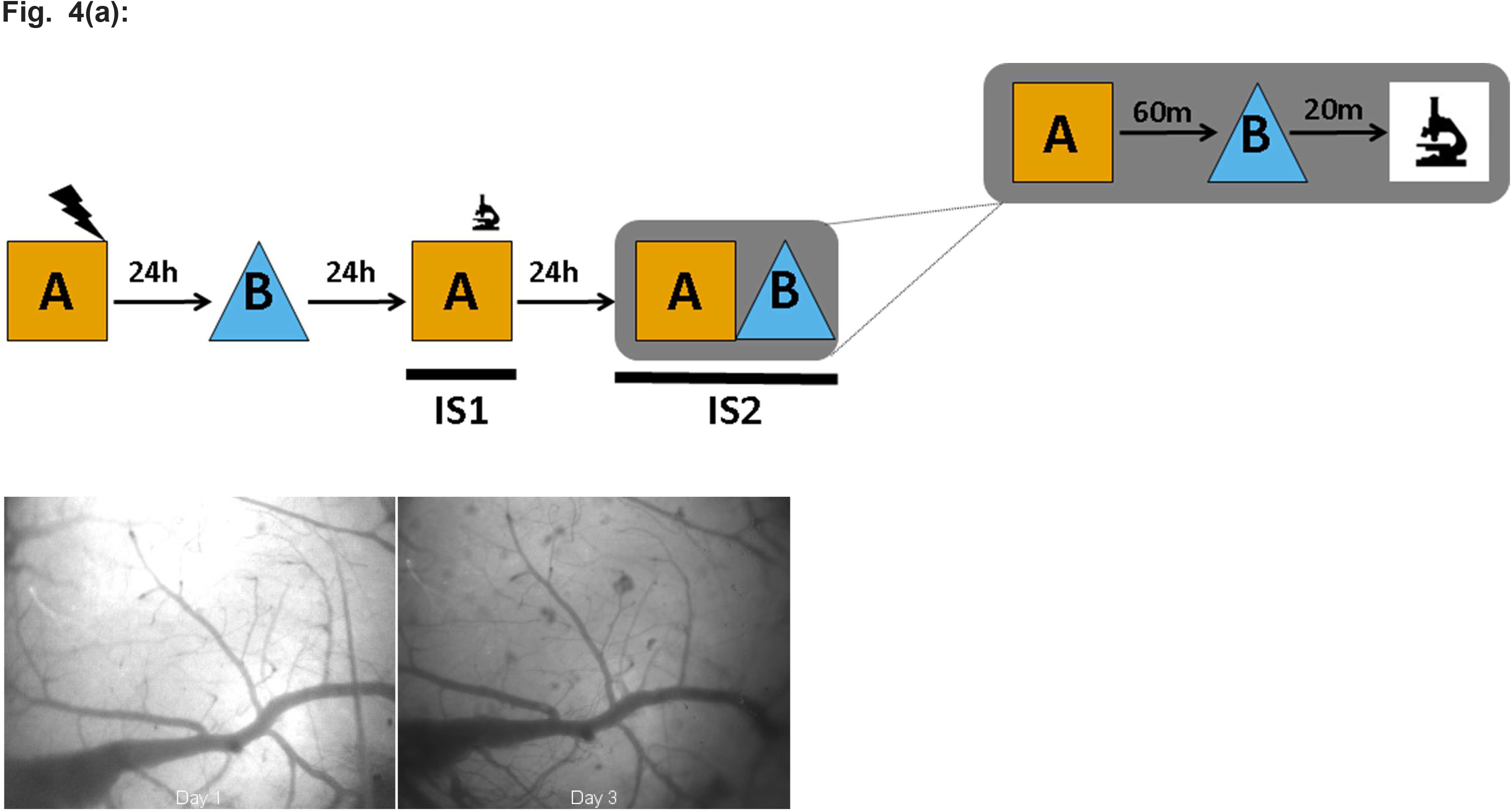
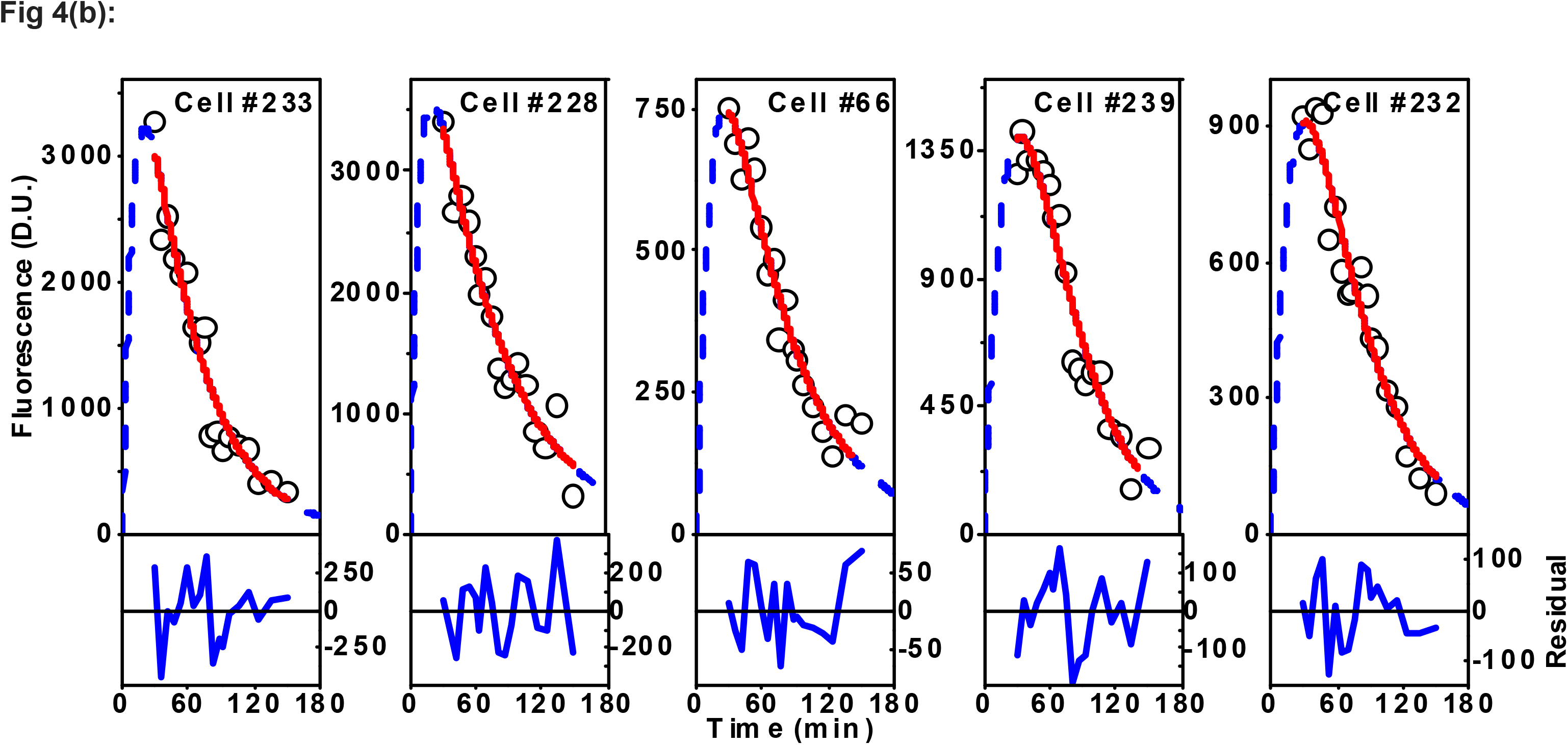
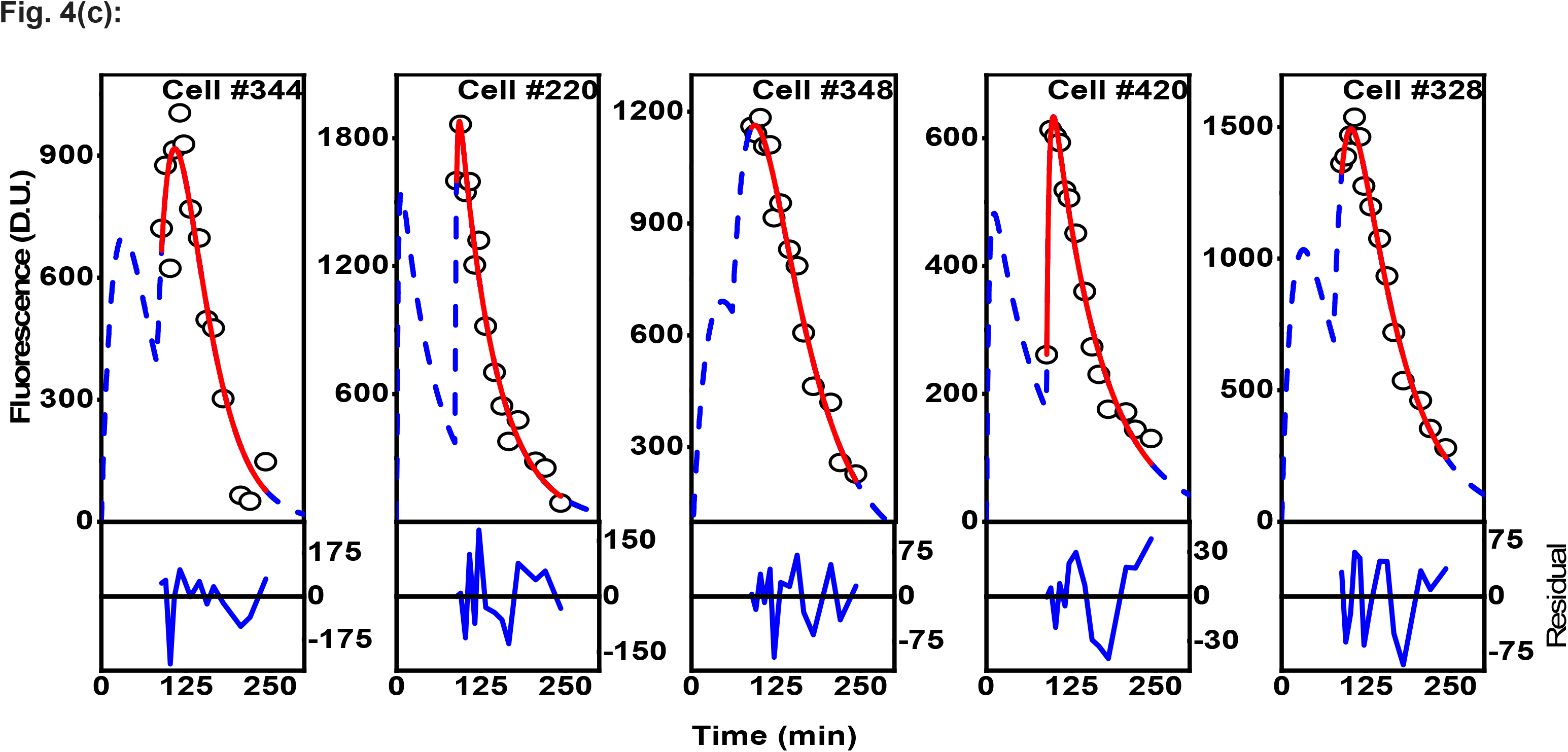
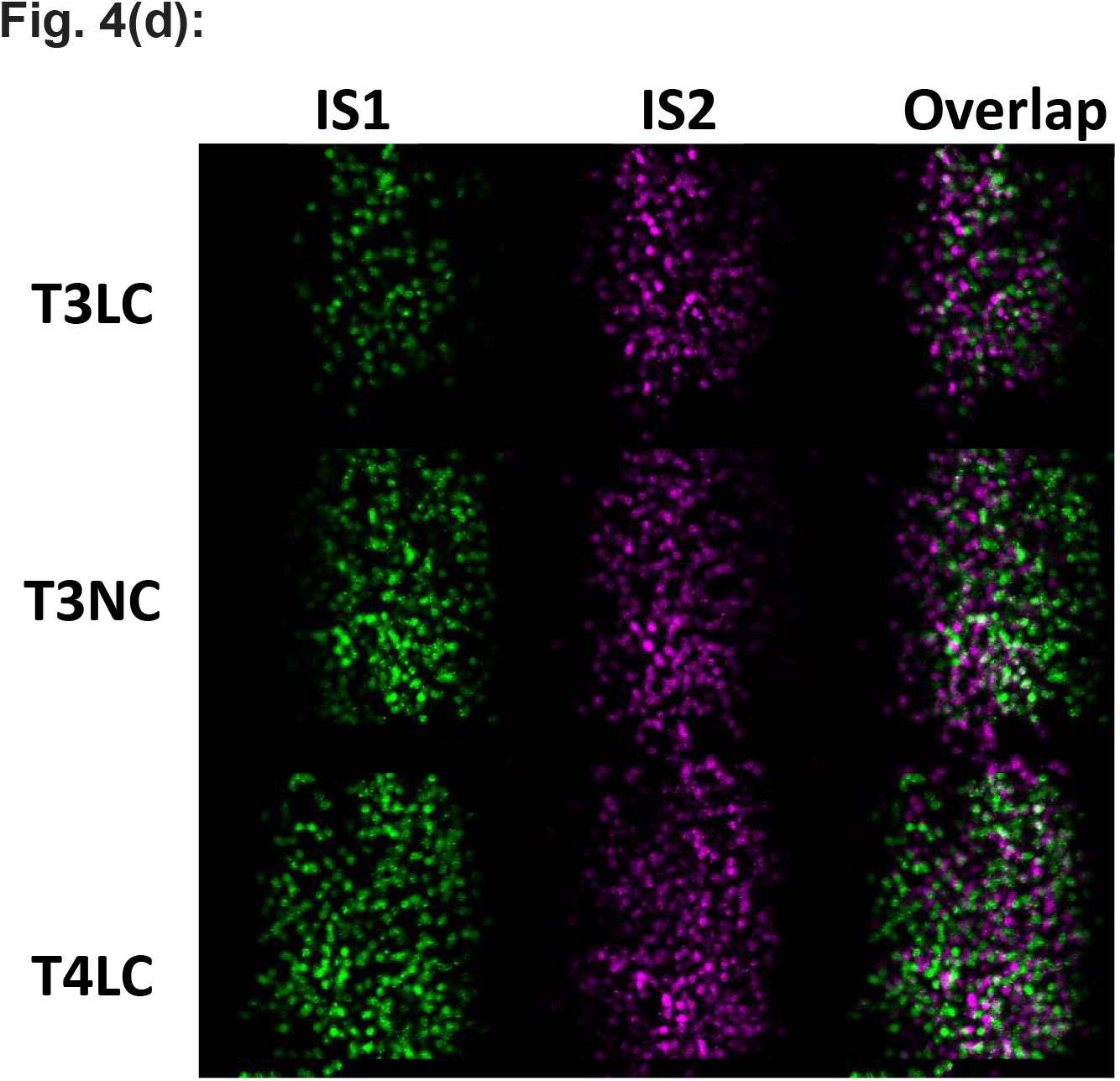
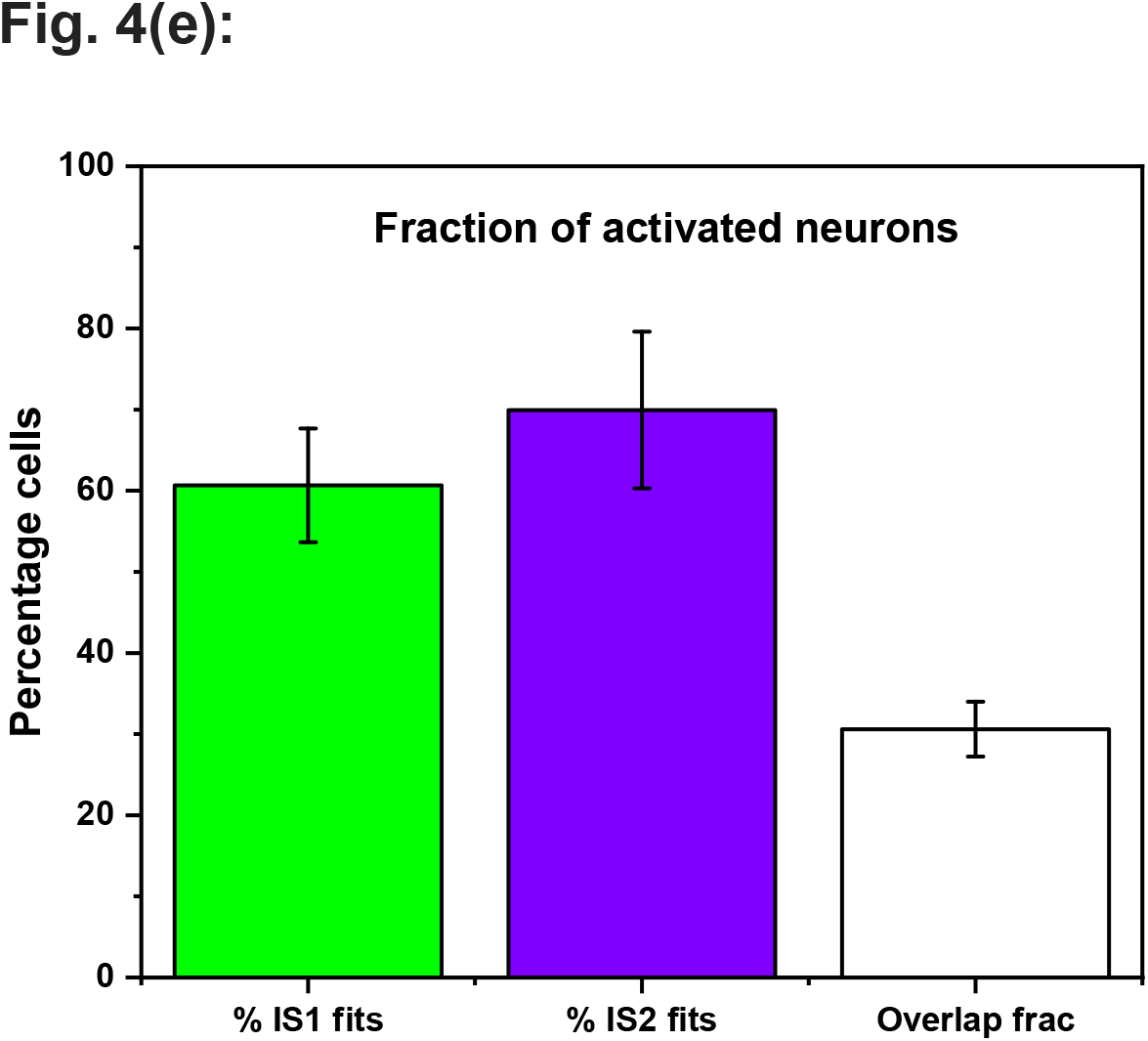
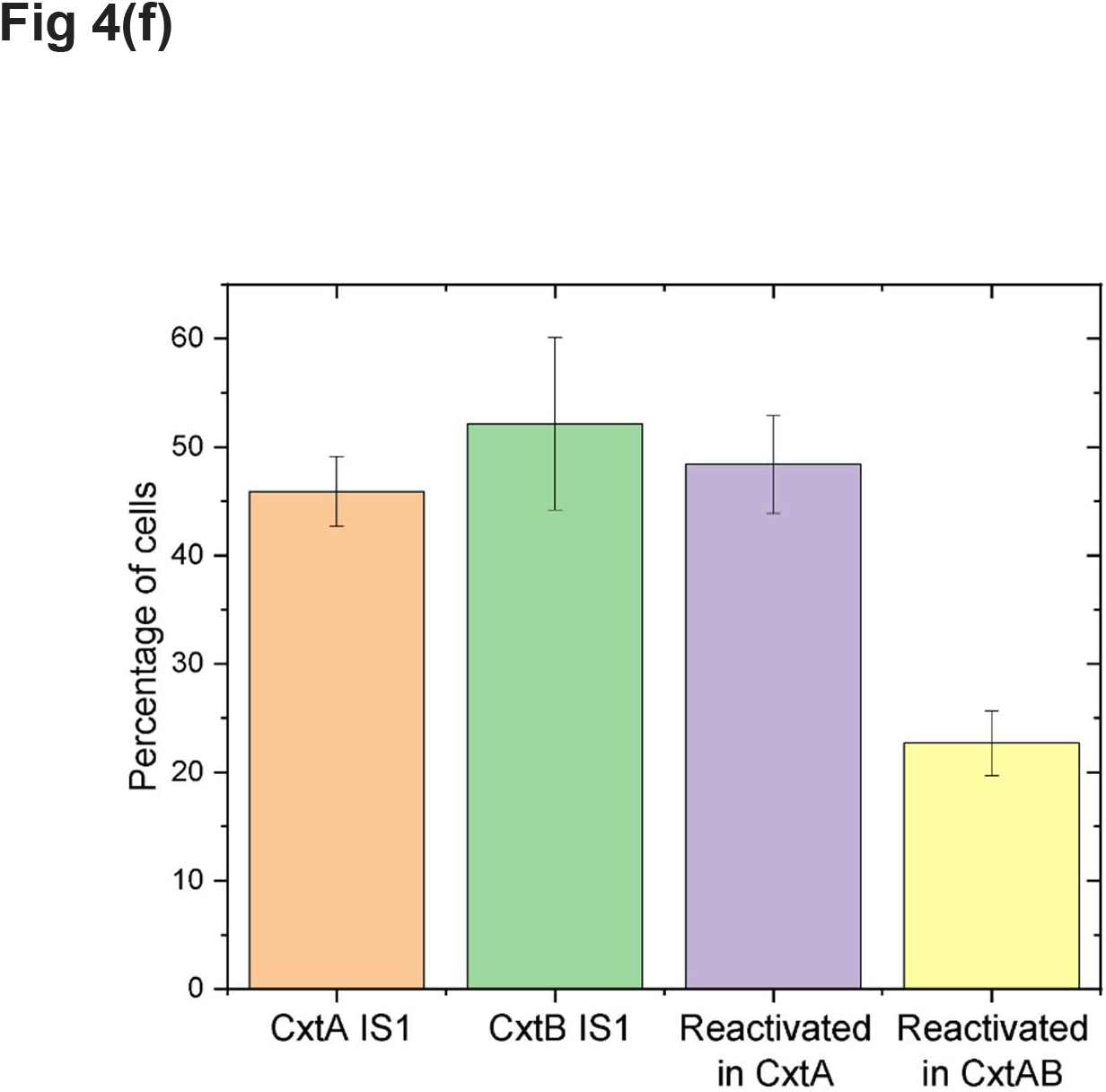
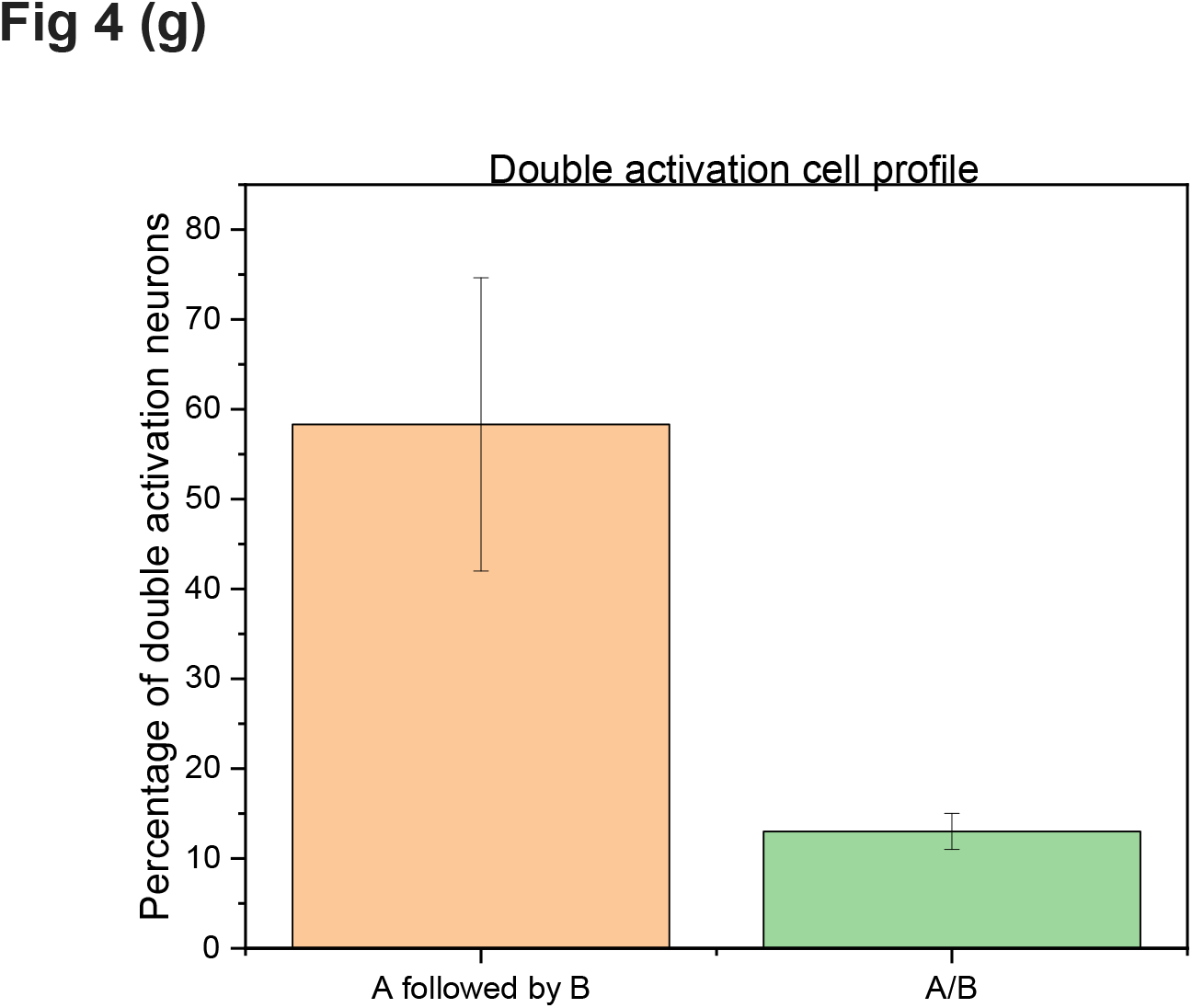
Identification and segregation of neuronal ensembles based on cfos protein temporal expression dynamics. (a) Schematic of behavioural paradigm used. (b) Representative plots of expression profile of single activation neurons. (c) Representative plots of expression profile of double activation neurons. The curve of the fit (red line) is explained better by DAC profile eq 2 rather than SAC profile eq 1. The dashed blue line shows the fit spanning the entire laboratory time frame of t = 0 mins where the mouse was exposed to the first context A, followed by t = 60 mins where the mouse was exposed to context B. (d) Representative snapshot of cfos neuronal ensembles at ~90mins on imaging sessions IS1(green) and IS2(magenta) with overlapping ensembles shown in white. (e) Fraction of activated neurons on days IS1, IS2 and their overlap. (f) Fraction of activated neurons on different retrieval events. Different bars represent activated neurons (SAC or DAC) in response to different cxt exposure - Orange: Cxt A on IS1, Green: CxtB on IS2, Purple: CxtA IS1 cells activated on CxtA IS2, Yellow: CxtA IS1 cells activated as DAC on CxtAB IS2. (g) The fraction of double activation cells in response to cxt A followed by cxt B on IS2 is significantly more than the fraction of double activation cells in response to single context exposure event

**Table 1:** Table summarizing the T_max_, k_f_ and k_d_ values from Gauss fit (in S2) to the respective distributions (Adj-R-Sq > 0.95).

| Transgenic mouse | $T_{max}$ (min) | $k_f$ ($\text{min}^{-1}$) | $k_d$ ($\text{min}^{-1}$) | |
| --- | --- | --- | --- | --- |
| cfos-shGFP | $41 \pm 1$ | $0.0369 \pm 4 \text{ E-4}$ | $0.0062 \pm 1\text{E-4}$ | $0.016 \pm 0.002$ |
| cfos-EGFP | $74.5 \pm 0.75$ | $0.0369 \pm 2\text{E-4}$ | $0.002 \pm 1\text{E-4}$ | $0.005 \pm 7\text{E-4}$ |

**Table 2:** Summary of fit parameters of data fit to equation 1.

| Fig No. | Cell No. | Amplitude<br>(D.U.) | Error in Amplitude | $k_f$<br>(min<sup>-1</sup>) | Error in $k_f$ | $K_d$<br>(min<sup>-1</sup>) | Error in $k_d$ | Adj-R-sq | AIC |
| --- | --- | --- | --- | --- | --- | --- | --- | --- | --- |
| Fig 2(d) | Cell #39 | 2640 | 39.68309 | 0.04605 | 0.00367 | 0.00176 | 1.41E-04 | 0.97354 | 85.43579 |
| Fig 2(d) | Cell #41 | 2576 | 37.18148 | 0.04616 | 0.00355 | 0.00168 | 1.34E-04 | 0.97265 | 83.87009 |
| Fig 2(d) | Cell #46 | 2262 | 32.49374 | 0.05358 | 0.00587 | 0.0011 | 1.30E-04 | 0.93118 | 84.81639 |
| Fig 2(d) | Cell # 05 | 2888 | 76.33026 | 0.04576 | 0.00633 | 0.00183 | 2.49E-04 | 0.92799 | 102.19985 |
| Fig 3(c) | Cell #28 | 541 | 26.39433 | 0.03675 | 0.00495 | 0.00514 | 5.31E-04 | 0.95524 | 97.85696 |
| Fig 3(c) | Cell #11 | 1050 | 148.97179 | 0.02585 | 0.00614 | 0.00895 | 0.00182 | 0.94341 | 123.23644 |
| Fig 3(c) | Cell #03 | 596 | 50.12666 | 0.02569 | 0.00406 | 0.00592 | 8.67E-04 | 0.93926 | 101.22503 |
| Fig 3(c) | Cell #38 | 461 | 27.40859 | 0.03473 | 0.0054 | 0.00487 | 6.20E-04 | 0.93042 | 98.06544 |
| Fig 3(e) | Cell #501 | 1183 | 148.68142 | 0.0593 | 0.02149 | 0.01404 | 0.00272 | 0.91592 | 150.73661 |
| Fig 3(e) | Cell #486 | 1089 | 104.53632 | 0.10002 | 0.07986 | 0.014 | 0.00265 | 0.86964 | 157.76146 |
| Fig 3(e) | Cell #475* | 3663 | 416.218 | 0.02647 | - | 0.02647 | 0.0036 | 0.87058 | 184.62851 |
| Fig 3(e) | Cell #430 | 1471 | 157.13412 | 0.07198 | 0.03514 | 0.0079 | 0.00176 | 0.79118 | 172.29792 |
| Fig 4(b) | Cell #233 | 4894 | 420.98677 | 0.09777 | 0.05748 | 0.02089 | 0.00332 | 0.9343 | 202.68672 |
| Fig 4(b) | Cell #228 | 4856 | 282.65906 | 0.11699 | 0.07415 | 0.01537 | 0.00164 | 0.94395 | 187.96317 |
| Fig 4(b) | Cell #66 | 1164 | 107.45433 | 0.07322 | 0.02521 | 0.017 | 0.00255 | 0.9444 | 146.82517 |
| Fig 4(b) | Cell #239* | 3807 | 261.832 | 0.03036 | - | 0.03037 | 0.00229 | 0.94465 | 174.71616 |
| Fig 4(b) | Cell #232 | 1802 | 492.2815 | 0.04495 | 0.0202 | 0.02197 | 0.00738 | 0.9378 | 159.5796 |

\*The $k_f$ and $k_d$ values for these cells were shared.

The fluorescence as a function of time shows a rapid increase in cfos-egfp protein that peaks at 74 + 0.7 minutes. Interestingly, previous studies report cfos protein expression peaks between 90-120 minutes for protein as detected by immunocytochemistry analysis (Barth et al., 2004a; Morgan et al., 1987). We argue that *in vivo* fluorescence signal is more sensitive in capturing cfos-egfp protein concentration level in a neuron in real time, compared to immunocytochemistry leading to the discrepancy in reported time range of maximal cfos protein expression. Given a good fit of the cFOS activation data, we asked if we could use such a model to predict expression of cFOS that occurs during memory formation.

### Behaviourally induced IEG protein expression described by the consecutive first-order kinetics expression indicate that the analytical equation generalises to different IEG – fluorophore constructs

Next, we investigated the expression kinetics of cfos in response to a behaviour using two different transgenic mice lines cFOS-EGFP and cFOS-shGFP. The mice were imaged following context recall and their time series data is collected as explained in the above section. Imaging time points ranged from 20 – 180 mins Fig 3(a). Fig 3 (c) shows the fit to data obtained from four neurons of cfos-egfp mice. Similarly, Fig 3(e) shows the fit of fluorescence obtained from neurons of cfos-shGFP mice. A total of 630 ROIs were identified for fluorescence extraction. These ROIs were used for data analysis the fit parameters describing the rate of formation(k_f_), decay(k_d_) and extent of activation (A0) were collected for individual cells for these mice and their distribution is compared in Supplementary Figure S2. During this process we observed that some of the cells had overlapping k_f_ and k_d_ values with high interdependency, we believe this is due to difference in when the cell got activated (start time) resulting lower density of data points for reliable estimating the k_f_, rather than the lack of fit. We have included some of these cell responses in Fig. 3e, 4b and 4c (Please see Table 2 and Table 3 footnote). Thus, to get reliable estimates of rate constants we obtain the histogram of these rate constants and fit to normal distribution to obtain the mean values. In case of decay constant k_d_ we see a bimodal distribution (verified through AIC) and we report both the decay constants in Table 1. Thus, we find the time to maximal activation estimated from Eq. 1 using the mean k_f_ and k_d_ values to be different for cFOS EGFP(74.5 ± 0.7 mins) and shGFP mice(41 ± 1 mins). This is in accordance with the genetic makeup of the mice and the properties of the transgene. In cFOS eGFP, the transgene is the fusion of EGFP and cFOS protein while in the cFOS shGFP just the GFP protein is expressed under cFOS promoter.

**Table 3:**
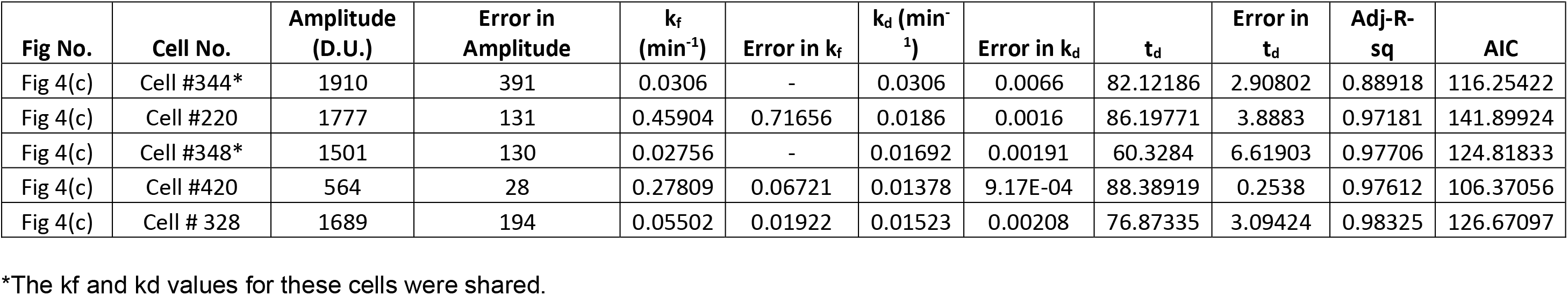
Summary of fit parameters of data fit to equation 2.

| Fig No. | Cell No. | Amplitude (D.U.) | Error in Amplitude | $k_f$ ($\text{min}^{-1}$) | Error in $k_f$ | $k_d$ ($\text{min}^{-1}$) | Error in $k_d$ | $t_d$ | Error in $t_d$ | Adj-R-sq | AIC |
| --- | --- | --- | --- | --- | --- | --- | --- | --- | --- | --- | --- |
| Fig 4(c) | Cell #344* | 1910 | 391 | 0.0306 | - | 0.0306 | 0.0066 | 82.12186 | 2.90802 | 0.88918 | 116.25422 |
| Fig 4(c) | Cell #220 | 1777 | 131 | 0.45904 | 0.71656 | 0.0186 | 0.0016 | 86.19771 | 3.8883 | 0.97181 | 141.89924 |
| Fig 4(c) | Cell #348* | 1501 | 130 | 0.02756 | - | 0.01692 | 0.00191 | 60.3284 | 6.61903 | 0.97706 | 124.81833 |
| Fig 4(c) | Cell #420 | 564 | 28 | 0.27809 | 0.06721 | 0.01378 | 9.17E-04 | 88.38919 | 0.2538 | 0.97612 | 106.37056 |
| Fig 4(c) | Cell # 328 | 1689 | 194 | 0.05502 | 0.01922 | 0.01523 | 0.00208 | 76.87335 | 3.09424 | 0.98325 | 126.67097 |

\*The $k_f$ and $k_d$ values for these cells were shared.

A good agreement of the experimental data with our model in both these mice lines following seizure as well as the behavioral activation suggests that our method can identify the neuronal ensemble that represents activation. Additionally, the fluorescence signal as a function of time is well described by the analytical expression irrespective of the transgenic mice used. Thus, it lends support to the hypothesis that the analytical equation can be generalised to the protein expression kinetics of other IEG (e.g., arc, zif).

### Identification and segregation of neuronal ensemble following single and dual context exposure

Using the established method of segregating differentially activated neurons, we sought to identify the neuronal ensembles that emerge in response to contextual fear conditioning learning. We used the following behavioural schematic (Fig 4 (a)) to train and image the contextual representation in cfos-shgfp transgenic mice. Briefly, the mice were trained to associate a foot shock in context A followed by safety training in context B the next day. Imaging sessions were carried out 24h after exposure to context B. First imaging session is that of exposure to context A after training (IS1 CxtA). 24 hours later, we image the mice following dual exposure to context A followed by context B. The exposures to contexts were 60 minutes apart. Thus, there were two imaging sessions: one following context A (IS1) and the other following context B on the second day of imaging (IS2). In IS1, the imaging session consisted of 20-180 minutes whereas in IS2 the imaging points consisted of 90-300 minutes, at intervals of ~10 minutes. Since there were ~2000 ROIs to be processed, analysed, and classified, we developed a software based workflow as described in Materials and Methods (Supplementary Figure S1). In both these imaging sessions, we see two kinds of cellular response: (I) cells that were activated once (single activation cells, SAC) and (II) twice (double activation cells, DAC) in response to context exposure. Fig 4(b) and 4(c) shows representative cfos expression of SAC and DAC fit to Eq. 1 and 2, respectively. The quality of the fit can be assessed through the residuals displayed at the bottom panel of each response. Table 2 and 3 summaries the fit parameters. Due to their difference in initial slope and curvature around the peak of DAC compared to SAC, DAC fit the double activation cell profile corresponding to Eq. 2 better as determined by our criteria (AIC, Adj r-sq >=0.5). Supplementary Figure S3 illustrates this behavior use an example cell response.The investigated ROIs fit either Eq. 1 or 2 on at least one of the imaging sessions for us to consider it a neuron. We find that 71 +6% of the investigated ROIs fall in this category. Fig 4(d) show the snapshot of the cellular activity at the RSc of cfos-shgfp transgenic mice. The green cells are the activated cells on imaging session 1 and the magenta are the activated cells on imaging session 2. The cells that got activated on both IS1 and IS2 is shown in the merge panel in white. We constructed these images from snapshots of the RSc at ~90 minutes after the first context exposure of the imaging session. This fraction though is not known so far in RSc, it is similar to what is observed in other regions of the brain (Vazdarjanova and Guzowski, 2004; Repa et al., 2001) we see an overlap of ~30% of neurons that were activated in response to contextual exposure on both imaging sessions (Fig 4(e) white bar).

Using our method, we segregate the activated neurons in this population into different groups:

1. neurons that were activated once after context A exposure in imaging session 1 (IS1 CxtA, Fig4(f) orange bar).
2. neurons that were activated once with its expression starting at t = 0 (when mouse was placed in context A chamber) after dual exposure, context A followed by B with a temporal separation of 60 mins in imaging session 2 (IS2 CxtA).
3. neurons that were activated once with its expression starting at t = 60 (when mouse was placed in context B chamber) after dual exposure, context A followed by B with a temporal separation of 60 mins in imaging session 2 (IS2 CxtB, Fig4(f) green bar).
4. neurons that were activated twice in response to context A exposure followed by context B in imaging session 2 (IS2 CxtAB)
5. Response of activated neurons that do not correspond to a behaviourally relevant event.

We see that the fraction of neurons activated in response to exposure to a single context for the first retrieval event is ~50% (Fig 4(f) CxtA orange (45.9+-3.2) and CxtB green (52.1+-7.9) bars) which is similar to what has been found in other regions of the brain with dense encoding(Vazdarjanova and Guzowski, 2004). We see responses of ~ 20% of neurons still shows cellular response profile that enables them to be classified as active in context A even though the exposure has occurred 90 mins prior to first imaging data point of imaging session 2. Of these activated neurons, ~50% of them where the neurons previously activated in Ctxt A (48.4±4.5, purple bar). Intriguingly, ~20% of the neurons activated in context B show expression dynamics consistent with dual activation profile, we interpret this as a neuronal response to context A as well as context B (22.6±2.9, yellow bar) exposure.

Next, we compare these groups with our behavioural scheme to reason and assign these different classes to the corresponding memory representations. Since IS1 was followed by retrieval event in context A and these cells responses are consistent with eq 1 representing single events, we assign them to context A cellular ensemble. Though, the activated cellular fraction of 49±3 % at RSc has not been reported before, it is comparable to the ensemble size typically obtained in other regions through conventional studies(Guzowski and Worley, 2001). Similarly, we assign the SAC from context B exposure and DAC from context A exposure of IS2 to context B cellular representation. We note this fraction 52±8% for context B is similar to that of context A cellular fraction. Thus, consistent with our hypothesis, these two contextual representations can be identified and assigned to different cell populations using cellular activation profiles. We argue that if context A cellular ensemble were to truly represent context A, a vast majority of the cells that were active in second session should have also been active in the first session. We estimated this fraction to be 48.4±4.5% (Fig 4(f) third bar), indicating that at least half the neurons that were activated in response to context A were repeatedly activated in response to context A. However, when we look at the fraction of DAC of IS2 that were activated previously in as SAC of IS1, we see only a fraction of 22.6 ± 2.9% (Fig 4(f) fourth bar). Since it was comparable to the chance activation (22.5%) we reason that these ensembles could be representing information other than context A. One of the defining events of IS2 is the arrival of the context B exposure after context A. It is known that contextual exposure could be temporally linked through cellular activation(Silva et al., 2009)(Cai et al., 2016). This being the case, we expect a larger fraction of activated cells following context A exposure in IS2 would be DACs. Thus, we compare the fraction of DACs seen in response to context B following context A exposure with that of DACs seen in response to a single context A or B (Fig 4 (g)). We see DACs of context AB (58.3 ±16.3%) to be significantly different (t test: p <0.002, one tailed) than DACs of either context A or B (~13 + 2%). Such an information is hard to obtain through conventional molecular probing methods since the comparisons are made at a population level instead of a single cell.

In summary, we can follow the IEG protein expression dynamics to identify and segregate the activation of neurons into different groups belonging to behavioural events. Conventionally, three probes are required to mark three retrieval events, however our method requires one IEG-fluorophore probe to mark all three retrieval events. Since our method utilizes the temporal profile of IEG expression, as opposed to intensity threshold, to classify neurons, we are able to increase the specificity with which we identify behaviourally relevant activated neurons.

Interestingly, contrary to conventional methods, we find cfos protein’s maximum expression takes place around ~74 mins for fos-egfp transgenic mice, and ~40 mins for fos-shgfp transgenic mice, instead of 90mins. We attribute this difference to our method’s ability to estimate cellular activation. In our method, we directly estimate the time to peak activation from individual cells whereas other methods are limited to estimating this by measuring the cell fraction rather than the cell activation (Wen et al., 2013; Barth et al., 2004b; Guzowski, 2002b). Such an estimate allowed us to probe a dual context exposure interaction and show that RSc indeed encodes this as described below.

For example, DAC in response to context A or context B only would be falsely identified as representing two contexts. More importantly, majority of the 1-ROI fit fraction, would have been classified as representing one of the contexts when in fact they do not represent IEG activation. While it is possible the lack of fit could represent a different model of activation profile, we see a vast fraction of these ROIs to be invariant with respect to time. Though their fluorescence is measured to be above the baseline. Further, we could identify the ensemble representations of different contextual memories in RSc as early as 24 hours following training. Strikingly, we could also identify a neuronal ensemble that possibly encodes the memory linkage when two contextual memories are presented close in time.

## Materials and Methods

### Transgenic mice

cFOS-EGFP (B6.Cg-Tg(Fos/EGFP)1-3Brth/J Stock no: 014135, Barth et al) and FOS-tTA(B6.Cg-Tg(Fos-tTA,Fos-EGFP*)1Mmay/J Stock no: 018306, Reijmers et al) transgenic mice were obtained from Jackson Laboratory, USA and maintained at the Central Animal Facility, IISc. All protocols were approved by the Institute Animal Ethics Committee.

### Craniotomy

Transgenic mice underwent a craniotomy to enable in vivo imaging (Vazdarjanova and Guzowski, 2004). A sterile 6mm converglass was positioned over the skull between the bregma and lambda, centred at retrosplenial cortex (RSc). The coordinates of the imaging area (RSc: 2mm from Bregma, 0.5mm laterally.) were arrived at by visualising the blood vasculature. The mice were anesthetized using a solution of fentanyl (0.05 mg/kg), midazolam (5 mg/kg), and medetomidin (0.5 mg/kg) dissolved in saline.

### Artificially and behaviourally induced IEG expression

Artificial IEG induction was produced by injecting mouse with 2mg/kg bicuculline intraperitonially to induce a seizure. Mild seizure symptoms were observed. On completion of the seizure, mouse was anesthetized with FMM to proceed to *in vivo* imaging.

Behaviour paradigm: Mice were trained to associate a mild foot-shock (0.7mA, 2s) in context A (70% ethanol, spaced grill floor) on training day (2 minutes 30 seconds, Day 1). The next day, these mice were placed in context B (20% ethyl acetate, smooth floor, triangle chamber feature) without shock for 2 mins 30 seconds (safety context training). After 24 hours (Day 3), mice were placed again in context A to test for memory recall. On Day 4, mice were placed in context A followed by context B separated by a time period of 60 minutes to test their ability to discriminate similar contexts at a recent time (24/48 hours).

### Imaging setup

In vivo imaging was performed on a custom-built two-photon setup based on a Zeiss upright microscope (Axio Examiner Z1) equipped with a 25× water immersion objective (NA 1.05, WD 2 mm, Olympus XLPLN25XWMP2). Femtosecond pulses from a ultrafast Ti:Sapphire laser (Newport, Tsunami) whose intensity was modulated using a half-wave plate (Thorlabs, AHWP05M) and a polarizer (Thorlabs, GL10-B) is as light source. The excitation beam is raster scanned using a galvo scanning mirrors before entering the microscope body and is focussed on the imaging plane using the objective lens. The fluorescence that is collected by the objective lens in an epi-illumination geometry is then separated using a dichroic before being detected by photomultiplier module (H7422, Hamatsu Corporation, Japan). The excitation wavelength of 900nm with power of 30mW at the objective was used to excite EGFP fluorophore. A galvanometric scanning X-Y mirror pair (Thorlabs, GVSM002) scanned the laser beam across the sample plane. The emitted fluorescence was collected and imaged onto a photomultiplier tube (Hamamatsu, H7422-40), driven by a power supply unit with temperature control (C8137-02). Emitted fluorescence signal was separated from excitation light using a dichroic mirror (Semrock, FF705-Di01), emission filters (Semrock FF01-520/70) and an aqueous solution of copper sulphate. The field of view of the imaging area was 500 microns by 500 μm.

A low noise current preamplifier (Stanford Research Systems, SR570) was used to amplify the photomultiplier tube photocurrent, which was further digitized using a data acquisition board (National Instruments, PCI-6110). ScanImage (r 3.8.1) software was used to interface instrument control and generation of galvometric scan command. Image acquisition was accomplished using a custom Matlab script interfaced with z-drive of the microscope. The digitized signal was analysed using Matlab, Origin and ImageJ for further analysis.

### Estimation of fluorescence from cFOS expression neurons

The in vivo images were analysed manually using a modified version of Time Series Analyzer plugin (Balaji 2014) https://imagej.nih.gov/ij/plugins/time-series.html in ImageJ to quantify the fluorescence signal. The modified version of the plugin is publicly available as Java Repository in GitHub (Meenakshi and Balaji 2020), (github link: https://github.com/TheNeurodynamicsLab/ImageJ_NDLPlugins). Supplementary Figure S1 describes the steps to extract the fluorescence signal from each neuron to obtain the fluorescence value of a neuron at a given time point. Briefly, for each individual neuron, the peak intensity at given time point was quantified by manually selecting the nucleus as the region of interest (ROI). The mean pixel intensity of the ROI through each z-stack was obtained and fit to a Gaussian function to estimate the activity of the ROI at a particular time point (Supplementary Figure S1).

### Classification of SACs and DACs through curve fitting

Curve fitting analysis of fluorescence as a function of time for each ROI was done in Origin(v2020b)’s user-defined NLFit function using Levenberg-Marquardt algorithm. The parameters were set as follows for data fitting to equation 1: Parameter A is initialised to the maximum fluorescence of the ROI observed in the imaging session, while rate constants k_f_ and k_d_ are initialsed to 0.01 and 0.001 respectively at the start of the Levenberg-Marquardt algorithm for least squares minimisation. For data fitting to equation 2, the additional parameter td was initialised with a value of 60 mins. Post data fitting to equation 1 and 2, the preferred model (SAC or DAC) was selected based on Akaike Information Criterion (AIC). In brief, AIC measures the information loss incurred in choosing a fit model given the observed data and degrees of freedom. Thus it takes into account the difference in number of parameters used in a fit as well as the goodness of the fit. A model with low AIC explains the observed data with minimal loss of information without over fitting, and hence is preferred. The goodness of fit was determined by an Adj-r-square value and and it was set to be greater than 0.5 to identify the selected ROI as an activated cell of the preferred model.

## Author Contributions

M.P, S.K and B.J. designed the experiments. M.P and S.K. collected the data from the behavioural and imaging experiments. M.P and B.J developed the theory, analysed the data and wrote the article.

## Acknowledgments

This work is funded by grants to:

BJ from SERB (EMR/2017/004155), DBT IISc Partnership, Ramanujan Fellowship, and Pratiksha Trust.

CSIR Fellowship to M.P (CSIR-09/079(2697)/2016-EMR-I) and CSIR Fellowship to S.K. (CSIR-09/079(2561)2012-EMR-I).

## Supplementary figures

**Supplementary Figure S1:**
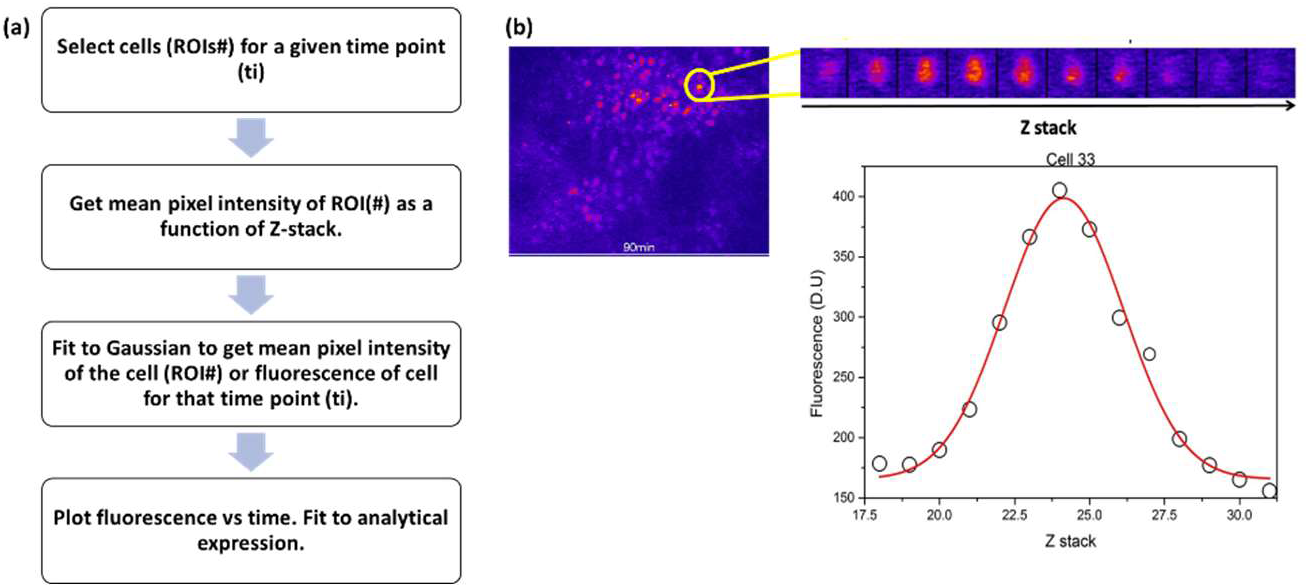
Extraction of fluorescence values. (a) Flowchart describing the steps for data extraction. (b) Left: Snapshot of field of view at 90 mins. Yellow circle and inset top represent one neuron. Top inset: Optical sections of the neuron at different Z positions in a stack. Bottom: Gaussian fit to obtain mean pixel intensity or fluorescence of a neuron at a given time point. The open circles represent the mean pixel intensity of the neuron at a slice/Z position. The red line represents the Gaussian fitting of mean pixel intensity as a function of Z position.

**Supplementary Figure S2:**
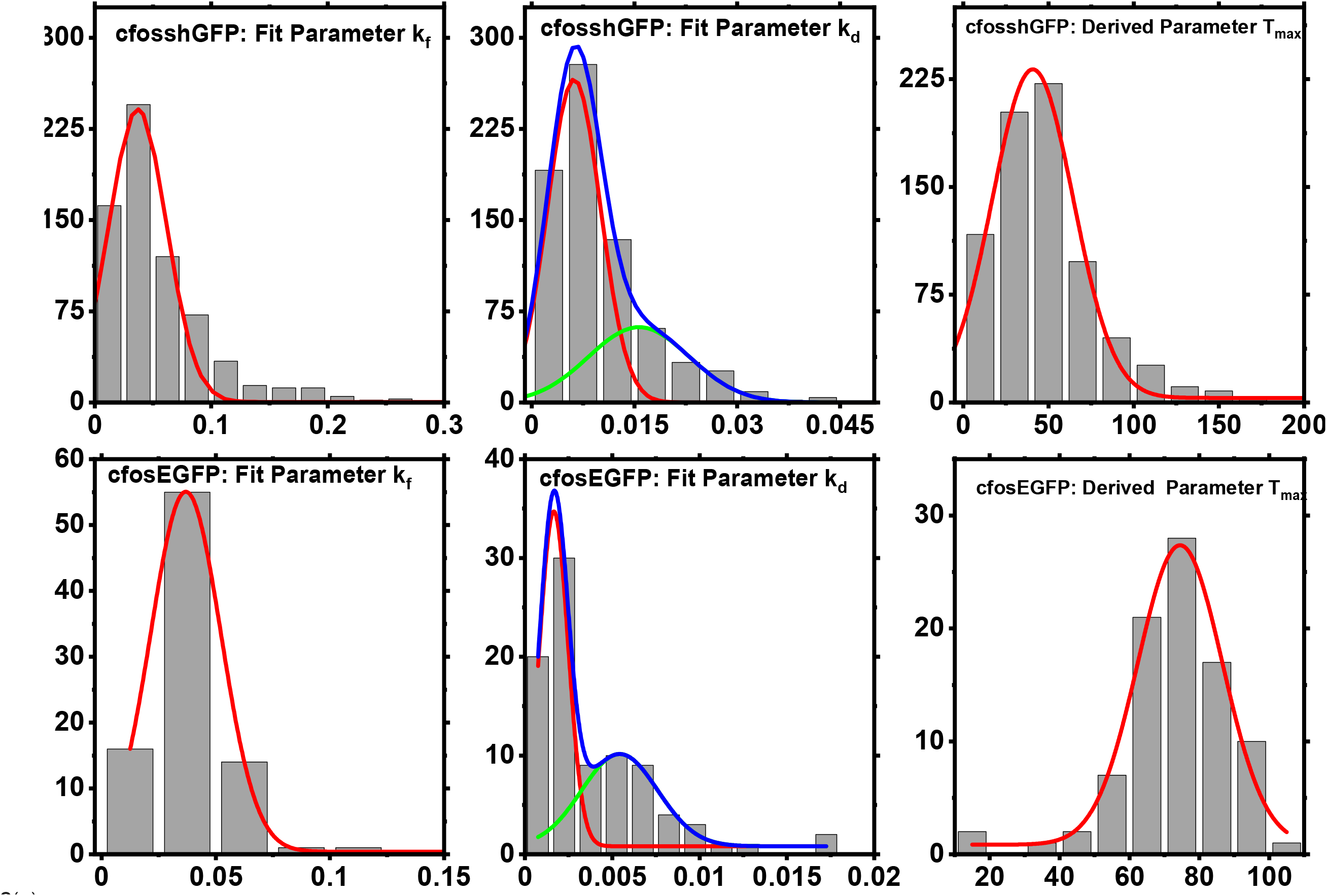
Frequency count distributions of fit and derived parameters: k_f_, k_d_ and T_max_ values. (a) Frequency count distributions were plotted (grey bars) using an appropriate bin width. The distribution was fit to a Gauss function (solid red line) to estimate the parameter value summarised in (b). Top row shows parameter distribution plots for cfos-shGFP transgenic mice (n = ~700 cells). Bins with counts >10 were considered for fit. We see a bimodal distribution of k_d_ values in cfos-shGFP mice indicating populations of neurons with different decay kinetics. Bottom row shows the corresponding plots for cfos-EGFP transgenic mice (n = ~90 cells).

**Supplementary Figure S3:**
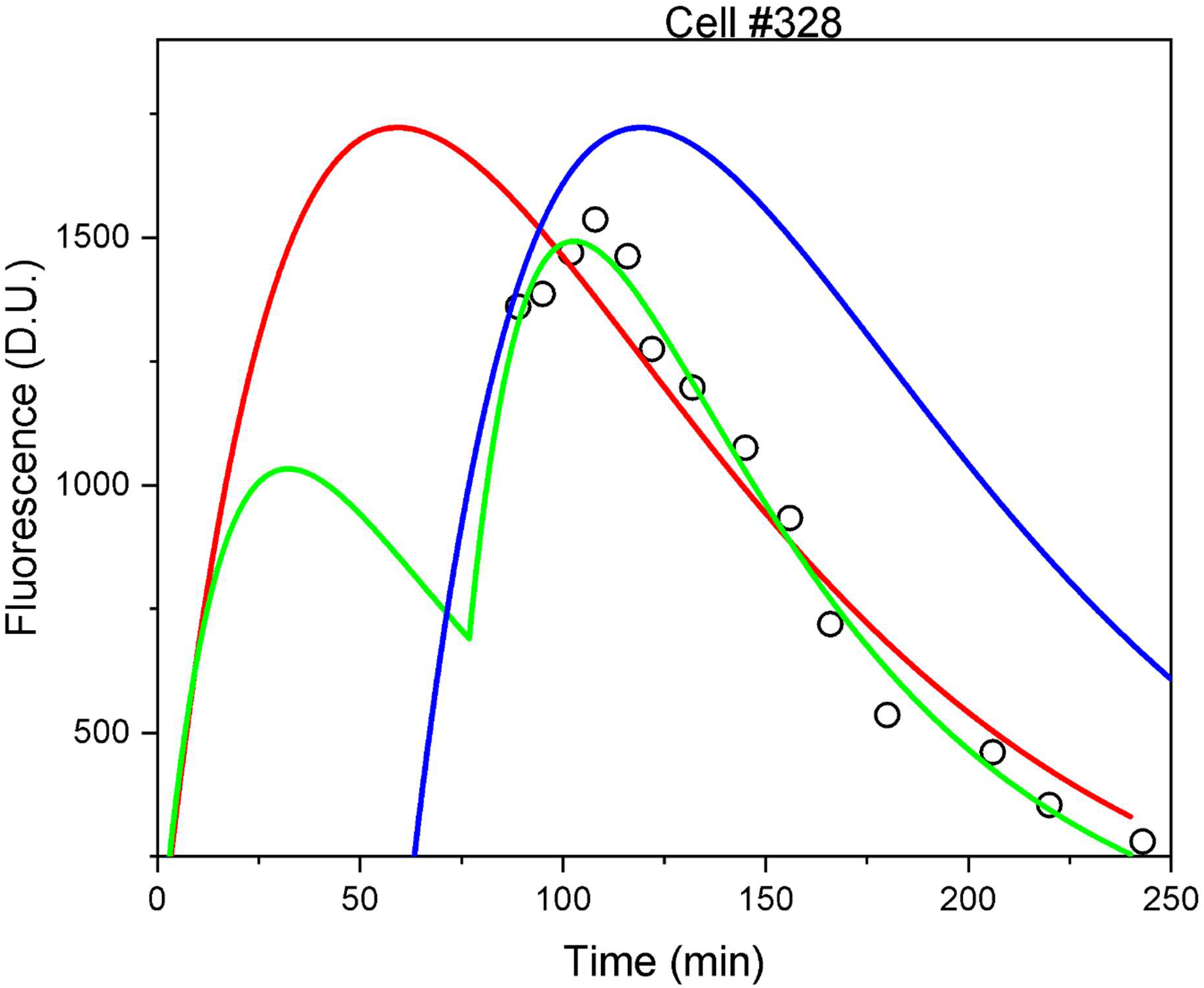
Comparison of a DAC fit to Eq 1 with (t = 60 min, solid green line) and without (t = 0 min, solid red line) delay along fit to Eq 2(solid blue line).

